# Human hindbrain organoids with segment-specific 5-HT neurons for motor modulation

**DOI:** 10.64898/2026.05.27.726154

**Authors:** Jinkui Zhu, Wei Pang, Yantong Liu, Yao Yin, Xiaoxiang Lu, Jianfeng Wei, Linlin Jiang, Wei Zhou, Yajing Cao, Qinzhi Zhang, Sisi Chen, Siyuan Chu, Zhaoxuan Ji, Die Hong, Yuanyuan Li, Yifei Tao, Xinrui Zhang, Chao Chen, Yangfei Xiang

**Affiliations:** School of Life Science and Technology, ShanghaiTech University, Shanghai 201210, China; MOE Key Laboratory of Rare Pediatric Diseases, School of Life Sciences, Central South University, Changsha 410083, Hunan, China; State Key Laboratory of Advanced Medical Materials and Devices, ShanghaiTech University, Shanghai 201210, China; Shanghai Clinical Research and Trial Center, Shanghai 201210, China

## Abstract

The serotonergic system regulates human mood, reward, and motor control, and its impairment contributes to debilitating psychiatric disorders. Serotonergic neurons (SNs) arise predominantly in the hindbrain and acquire segment-specific identities and functions; however, the lack of human models remains a challenge to understanding the human serotonergic system. Here, we generate human serotonergic-enriched rostral and caudal hindbrain organoids (hsrHOs and hscHOs) that produce segment-specific SNs with regional identities corresponding to rhombomeres r2–r3 and r5–r8, respectively. hsrHOs and hscHOs produce SNs that recapitulate their *in vivo* counterparts in transcriptomic profiles, function, and ascending or descending projection preferences. Assembly with human neuromusculoskeletal organoids enables modeling of serotonergic modulation of motor control. This system further reveals that deficiency of the schizophrenia risk gene DISC1 disrupts human serotonergic development and impairs motor modulation. Together, our study establishes a human organoid platform for modeling development, circuit function and disease-associated dysfunction of the serotonergic modulatory system.

## Introduction

The serotonergic system is broadly involved in the regulation of affect, reward processing and the coordination of motor control^1-3^. Within the central nervous system (CNS), serotonin (5-hydroxytryptamine, 5-HT) is produced predominantly by neuronal populations in the hindbrain raphe nuclei group (Fig. 1a), whose SNs have subsequently been recognized as a heterogeneous system^4,5^. Distinct SNs subtypes exhibit different molecular identities and projection patterns, thereby subserving specialized functions^6-9^. Dysfunctions of the serotonergic system has contributed to psychiatric disorders, including schizophrenia and major depressive disorder^10^. However, how SNs diversity arises, and how perturbations in this system contribute to disease pathogenesis is still poorly understood. Moreover, conventional rodent models have been questioned because of developmental differences relative to human SNs^11^. Therefore, approaches that can faithfully recapitulate the diversity of human SNs are essential for dissecting their development and for advancing our understanding of related disorders.

**Fig. 1.**
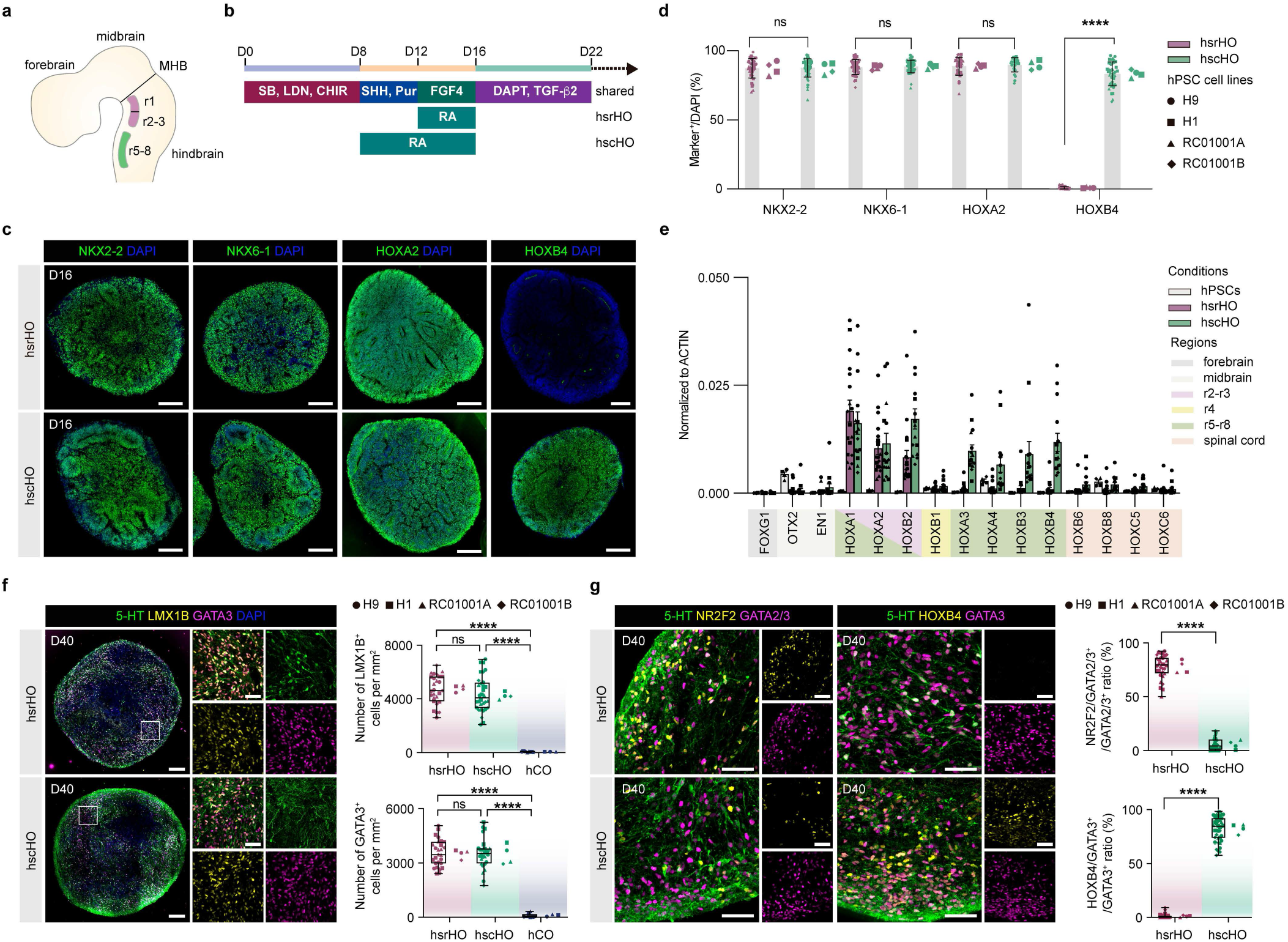
Generation of hsrHOs and hscHOs from hPSCs. **a**, Schematic diagram showing the SNs arising from different rhombomeres *in vivo*. MHB, midbrain-hindbrain boundary; r1, rhombomere 1; r2-3, rhombomere 2-3; r5-8, rhombomere 5-8. **b**, Differentiation conditions for hsrHOs and hscHOs. SB, SB431542; LDN, LDN193189; CHIR, CHIR99021; SHH, Sonic Hedgehog; Pur, purmorphamine; FGF4, fibroblast growth factor 4; RA, retinoic acid; TGF-β2, transforming growth factor beta 2. **c**, Representative immunostaining images of NKX2-2, NKX6-1, HOXA2 and HOXB4 of day 16 hsrHOs and hscHOs generated from H9 hESC line. **d**, Quantification of NKX2-2-, NKX6-1-, HOXA2- and HOXB4-positive cells of day 16 hsrHOs and hscHOs (NKX2-2, n = 39 of hsrHOs and n = 51 of hscHOs derived from 4 hPSC cell lines, four to eight differentiations for each cell line, p = 0.9145, two-tailed Mann-Whitney test; NKX6-1, n = 42 of hsrHOs and n = 48 of hscHOs derived from 4 hPSC cell lines, five to eight differentiations for each cell line, p = 0.7067, two-tailed Mann-Whitney test; HOXA2, n = 32 of hsrHOs and n = 30 of hscHOs derived from 4 hPSC cell lines, four to six differentiations for each cell line, p = 0.5998, two-tailed Mann-Whitney test; HOXB4, n = 53 of hsrHOs and n = 43 of hscHOs derived from 4 hPSC cell lines, four to eight differentiations for each cell line, ****p < 0.0001, two-tailed Mann-Whitney test). **e**, Relative gene expression by qPCR at day 22 of hsrHOs and hscHOs of rostrocaudal axis relative genes (hPSCs, n = 4 hPSC cell lines; hsrHOs, n = 18 differentiations from four hPSC cell lines; hsrHOs, n = 15 differentiations from four hPSC cell lines). **f**, Representative immunostaining images of SNs and quantification of LMX1B or GATA3 positive cells of day 40 hsrHOs, hscHOs and hCOs (LMX1B, n = 29 of hsrHOs and n = 30 of hscHOs derived from 4 hPSC cell lines, four to seven differentiations, n = 9 of hCOs derived from 3 hPSC cell lines, three differentiations, p = 0.0811 for hscHO versus hsrHO,****p < 0.0001 for hCO versus hsrHO, ****p < 0.0001 for hCO versus hscHO, two-tailed Welch’s t test; GATA3, n = 27 sections of hsrHOs and n = 28 sections of hscHOs derived from 4 hPSC cell lines, four to seven differentiations, n = 9 sections of hCOs derived from 3 hPSC cell lines, three differentiations, p = 0.8468 for hscHO versus hsrHO,****p < 0.0001 for hCO versus hsrHO, ****p < 0.0001 for hCO versus hscHO, two-tailed Welch’s t test). **g**, Representative immunostaining images and quantification of NR2F2 and GATA2/3 double-positive SNs or HOXB4 and GATA3 double-positive SNs of hsrHOs and hscHOs (For NR2F2, n = 32 of hsrHOs and n = 36 of hscHOs derived from 4 hPSC cell lines, three to five differentiations, ****p < 0.0001, two-tailed Mann-Whitney test; For HOXB4, n = 39 of hsrHOs and n = 39 of hscHOs derived from 4 hPSC cell lines, three to five differentiations, ****p < 0.0001, two-tailed Mann-Whitney test). Scale bars, 200 μm (c, f), 50 μm (f(zoom), g).

In general, the segmentation of the hindbrain into rhombomeres provides a rostrocaudal coordinate system for neuronal specification, while SHH-dependent ventral patterning is required for SN fate acquisition^12,13^. SNs arising from r1-r3 contribute to the rostral group, whereas those derived from r5-r8 populate the caudal group^5,14^. This segmental developmental origin provides a framework for understanding the diversity in serotonergic neuronal system. In recent years, the differentiation of region-specific brain organoids from human pluripotent stem cells (hPSCs) has provided a major opportunity to investigate the development of inaccessible brain regions and to model associated disorders^15-21^. Although two-dimensional (2D) differentiation protocols for SNs from hPSCs have been reported^22,23^, robust three-dimensional (3D) strategies for generating rhombomere-patterned hindbrain organoids enriched in regionally specified SNs remain lacking. Overcoming the inherent limitations of 2D systems would enable these neurons to better recapitulate interactions with other tissues, thereby make it possible to model multisystem crosstalk and disease modeling^15,24-27^. We previously developed approaches to model human thalamic reticular nucleus and medullary SpV nucleus^28,29^, indicating the feasibility of precisely recapitulating subregional brain structures through the modulation of relevant signaling pathways. Here, we developed a protocol for generating rhombomere-patterned hindbrain organoids enriched in segment-specific SNs. By fine-tuning retinoic acid (RA) treatment, we generated human hindbrain organoids that recapitulate the identities of the rostral hindbrain rhombomeres r2–r3 and the caudal hindbrain rhombomeres r5–r8, respectively. Notably, SNs generated under these two conditions acquired distinct segment-specific identities, closely resembling their *in vivo* counterparts in both transcriptomic profiles and projection patterns, and the protocol was robustly reproducible across multiple hESC and hiPSC lines. We refer to these two organoids as human serotonergic-enriched rostral hindbrain organoids (hsrHO) and human serotonergic-enriched caudal hindbrain organoids (hscHO), respectively. SNs in hsrHOs and hscHOs underwent maturation, released 5-HT, and responded to selective serotonin reuptake inhibitor (SSRI) treatment, while exhibiting distinct ascending or descending projection preferences. When fused with human neuromusculoskeletal organoids (hNMSOs) that we previously established^25^, these assembloids, termed human hindbrain–spinal–muscular assembloids (hHSMAs), recapitulated the modulatory effects of SNs on motor control. Using this system, knockout of the schizophrenia risk gene DISC1 (Disrupted in Schizophrenia 1) revealed morphological, electrophysiological and impaired motor control, offering insights into how serotonergic dysfunction may contribute to schizophrenia pathogenesis.

### Generation of hsrHOs and hscHOs from hPSCs

The hsrHOs and hscHOs were generated based on a static-to-spinning culture strategy we previously established^15-17,24,25,28-30^. During neuroectoderm specification induced by dual SMAD inhibition, we activated Wnt signaling, followed by sonic hedgehog (SHH) signaling, to induce ventral hindbrain identity. Temporal modulation of RA exposure was used to specify rhombomeres-specific progenitor fates. Fibroblast growth factor 4 (FGF4) was applied, followed by transforming growth factor-β2 (TGF-β2), as both factors have been reported to be critical for driving progenitors toward a serotonergic neuronal fate^22,31^. NOTCH signaling was blocked using DAPT to promote organoid maturation. Under this strategy (Fig. 1b), early-stage hsrHOs and hscHOs (day 16) were enriched with NKX2-2 and NKX6-1 positive ventral progenitor cells but not PAX7 positive dorsal progenitor cells (Fig. 1c, d and Extended Data Fig. 1a-e). Prolonged RA treatment (day 8 to day 16) enhanced caudalization, leading to the production of cells positive for HOXB4, a marker of hindbrain rhombomeres r5–r8. In contrast, short-term RA treatment (day 12 to day 16) led to the production of cells positive for HOXA2 but not HOXB4, consistent with the r2-r3 identity (Fig. 1c, d). Gene expression analysis at day 22 further showed that hsrHOs and hscHOs expressed distinct rostrocaudal marker genes corresponding to rhombomeres r2–r3 and r5–r8, respectively, rather than markers of forebrain, midbrain or spinal cord identity. (Fig. 1e). We also confirmed that four different hPSC lines (hESC lines: H9 and H1; hiPSC lines: RC01001A and RC01001B) could establish similar rostral-caudal identities during hsrHOs and hscHOs differentiation (Fig. 1d, e and Extended Data Fig. 1b-e). By day 22, immunostaining revealed the expression of serotonergic progenitor markers (LMX1B and GATA3), as well as 5-HT^+^ SNs at the periphery of the organoids (Extended Data Fig. 1f, g).

By day 40, hsrHOs and hscHOs were near-homogenously distributed with SNs labeled by 5-HT, LMX1B, and GATA3, distinct from our previously established human cortical organoids (hCOs) ^15^ (Fig. 1f and Extended Data Fig. 2a). Notably, SNs generated in hsrHOs and hscHOs acquired distinct regional identities. Specifically, in hscHOs, the majority of SNs were HOXB4-positive (81.86% ± 11.58), whereas in hsrHOs, the majority were NR2F2-positive (77.94% ± 11.02) (Fig. 1g), consistent with previous in *vivo* findings^6,7^ of their gene expression diversity. Besides hsrHOs and hscHOs derived from H9 hESCs, organoids derived from H1 hESCs, and hiPSC lines RC01001A and RC01001B, all showed similar lineage development and regional pattern (Extended Data Fig. 2b-d and Fig. 1g), demonstrating the reproducibility of the protocol across multiple hPSC lines. By day 40, immunostaining revealed tryptophan hydroxylase 2 (TPH2), the rate-limiting enzyme for 5-HT biosynthesis and a marker of mature SNs, expressed in a subset of SNs in both hsrHOs and hscHOs (Extended Data Fig. 2e, f).

Since several neuronal types arise from ventral NKX2-2-positive progenitors, we dissociated day 40 hsrHOs and hscHOs, reseeded the cells, and performed immunostaining to examine the neuron types present in organoids. We identified four distinct neuronal types: serotonergic (5-HT^+^) neurons, visceromotor (TH^+^) neurons, excitatory (vGlut2^+^) neurons, and inhibitory (GAD1^+^) neurons, with the serotonergic population constituting the predominant majority in both hsrHOs (56.92% ± 8.90) and hscHOs (52.40% ± 8.39) (Extended Data Fig. 3a-e). Quantitative analysis showed that the neuron types and their proportions were consistently established across batches derived from the four hPSC cell lines (Extended Data Fig. 3e).

It is known that the initial cell seeding density may affect the organoid patterning^32,33^. We found that reducing the initial seeding density from 500 to 100 cells (H9 hESCs) caused a posterior shift along the rostro-caudal axis without altering dorsal-ventral patterning: hscHOs, which previously exhibited an r5–r8 identity, instead expressed the spinal cord marker HOXC9, whereas hsrHOs, which originally corresponded to r2–r3, shifted to express the r5–r8 marker HOXB4 (Extended Data Fig. 4a, b). To facilitate reproducibility across different cell lines, we monitored organoid diameter across day 2 to day 40 and found that hsrHOs and hscHOs derived from all four hPSC lines converged to a similar size of approximately 800 µm in diameter by day 16 (Extended Data Fig. 4c). We therefore recommend using the day 16 diameter as a reference and, when necessary, combining it with gene expression and immunostaining analysis to determine the appropriate initial seeding cell number for newly introduced cell lines.

While RA treatment established the rostrocaudal identities of hsrHOs and hscHOs, we found that combined activation of SHH and FGF4 signaling was essential for SN differentiation. Specifically, we performed differentiation in which the timing and/or dosage of SHH and FGF4 supplementation was varied (Extended Data Fig. 4d). In general, prolonged SHH activation significantly upregulated the expression of floor plate associated genes^6^, whereas SHH withdrawal led to a loss of ventral progenitors; omission of FGF4 markedly reduced the expression of SN-associated markers (Extended Data Fig. 4e-g). Thus, appropriate SHH-mediated ventralization together with FGF4 exposure is essential for the generation of SNs.

### Segment-specific molecular features of hsrHOs and hscHOs

To further characterize lineage development, we used single-cell RNA sequencing (scRNA-seq) to profile cell composition of day 40 hsrHOs and hscHOs derived from H9 hESCs. The hsrHOs and hscHOs profiles were integrated using canonical correlation analysis (CCA) and visualized in a UMAP embedding. Eight major cell types were identified, including a small number of floor plate cell (FOXA2^+^, SHH^+^), neural progenitor cell (NPC; SOX2^+^, NES^+^), immature neuron (ASCL1^+^), neuron. Neural clusters included SN (FEV^+^, GATA3^+^), visceral motor neuron (CHAT^+^, PHOX2B^+^), glutamatergic neuron (SLC17A6^+^) and GABAergic neuron (GAD1^+^) (Fig. 2a, b and Extended Data Fig. 5a, b). Consistent with our immunostaining-based quantification, SNs comprised the largest neuronal population (Extended Data Fig. 5b). Interestingly, scRNA-seq and immunostaining also revealed a population of oligodendrocyte progenitor cells (OPC; CSPG4^+^, SOX10^+^) (Fig. 2a, b and Extended Data Fig. 5c, d), which give rise to myelinating oligodendrocytes in the brain. We next identified significantly up-regulated genes in SNs compared to all other cells and performed Gene Ontology (GO) analysis. Notably, GO terms associated with SNs function (e.g., serotonin uptake, metabolism and transport), as well as neuronal developmental processes (e.g., axonogenesis) were significantly enriched (Fig. 2c). Next, we mapped all neural clusters to E13.5 mouse brain ISH data using VoxHunt and found that neurons in hsrHOs most closely resembled a pontine identity, whereas neurons in hscHOs most resembled a medullary identity (Fig. 2d). RNA velocity analysis revealed a progenitor to neuron trajectory and SNs related genes (e.g., FEV, LMX1B and TPH2) were highly expressed at late stages along pseudotime (Extended Data Fig. 5e-g). To further characterize the molecular features of SNs, we extracted all SNs and classified them as mature (GATA3^+^, TPH2^+^) or immature (GATA3^+^, TPH2^-^) based on TPH2 expression (Fig. 2e and Extended Data Fig. 5j). We then mapped these neurons onto a primary human SNs reference atlas using an anchor-based label transfer methods (Fig. 2f and Extended Data Fig. 5h-j). We found that immature and mature SNs in hsrHOs showed the highest correlation with pontine SNs, whereas those in hscHOs showed the highest correlation with medullary SNs, when compared with primary brain tissues. Notably, a set of genes highly expressed in human pontine or medullary SNs was also detected in hsrHOs or hscHOs, respectively (Fig. 2g, h). These genes include those previously identified in *vivo* markers distinguishing SNs identities across regional subtypes, such as NR2F2, NR2F1, MEIS2 and HOXB3, as well as additional genes not previously highlighted, including SLC35F3, ZNF804B, SHOX2 and ID2 (Fig. 2i). These results suggest that hsrHOs and hscHOs preserve *in vivo* region-specific transcriptomic features, supporting their utility as a robust platform for studying human SNs diversity.

**Fig. 2.**
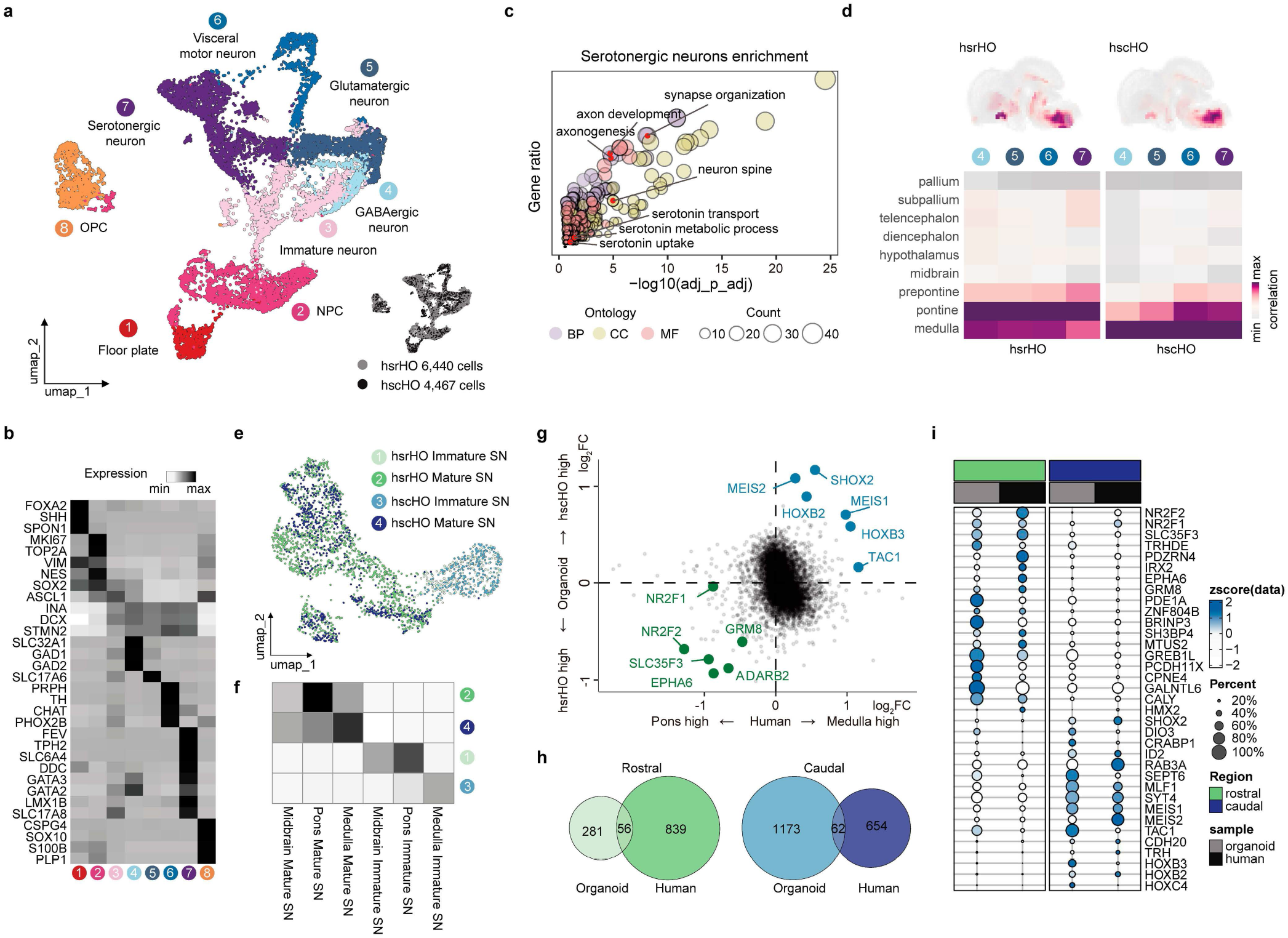
Molecular features of hsrHOs and hscHOs. **a**, UMAP embedding of integrated hsrHOs and hscHOs scRNA-seq profile, colored by annotated cell type or sample. **b**, Expression of representative marker genes in scRNA-seq profile. **c**, Gene Ontology analysis of up-regulated genes in SN clusters compared to all other cells. **d**, Voxhunt analysis showing the similarity of neural cells in hsrHOs and hscHOs to developing mouse brain (embryonic day 13.5). **e**, UMAP embedding of subcluster of SNs in hsrHOs and hscHOs. **f**, Heatmap showing the transcriptome similarity of SNs in hsrHOs and hscHOs to human SN reference. **g**, Log_2_(fold change) between rostral and caudal SNs showing the consistency between *in vitro* organoid-derived SNs and SNs from human brain tissue. **h**, Venn plots showing the overlaps of rostral-caudal SNs differentially expressed genes detected in organoids and human brain tissue. **i**, Dot plot showing the expression patterns of differentially expressed genes between rostral and caudal SNs across organoids and human brain tissue.

We next used bulk RNA-seq to assess the reproducibility of transcriptomic features in day 40 hsrHOs and hscHOs across four hPSC lines and three experimental batches per line (Extended Data Fig. 6a). In general, samples clustered by differentiation condition rather than by cell line (Extended Data Fig. 6b, c). Replicates of both hsrHOs and hscHOs exhibited a ventral hindbrain-specific identity, characterized by expression of ventral markers (NKX2-2 and NKX6-1) and hindbrain HOX genes, but not dorsal markers (OLIG3 and PAX7), hPSC markers, forebrain markers, midbrain markers, or spinal cord markers (Extended Data Fig. 6d). Bulk transcriptome deconvolution identified the major cell clusters detected by scRNA-seq in corresponding conditions, and the inferred cell fractions including SNs and OPCs were consistent across replicates and across cell lines (Extended Data Fig. 6e). Differentially expressed genes (DEGs) between hsrHOs and hscHOs were associated with regional identity and highly consistent across cell lines (Extended Data Fig. 6f, g). Overall, these data show that our differentiation protocol robustly established distinct rostral and caudal hindbrain identities and generated segment-specific SNs, with consistency across cell lines and batches.

### Functional development of hsrHOs and hscHOs

To assess long-term growth and functional characteristics of hsrHOs and hscHOs, we prepared 300 µm sections of day 30 organoids and cultured them at an air-liquid interface. By day 80, abundant GFAP-positive astrocytes were present in both hsrHOs and hscHOs (Fig. 3a). In addition, consistent with the emergence of OPCs at day 40 (Extended Data Fig. 5c, d, 6e), MBP-positive oligodendrocytes were also detected in both conditions (Fig. 3a). These glial cell populations are critical for neuronal survival and function, and notably, the generation of oligodendrocytes has been difficult to achieve in brain organoid models^34-36^. As the rate-limiting enzyme in 5-HT biosynthesis, TPH2 serves as a definitive marker of serotonergic identity and functional maturity^37,38^. We found that the proportion of TPH2-positive SNs was significantly increased at day 80 compared with day 40 in both hsrHOs and hscHOs (Fig. 3b), indicating progressive serotonergic development during long-term culture. To access electrophysiological activity of hsrHOs and hscHOs, we performed multi-electrode array (MEA) recording on day 90 organoid sections. Both hsrHOs and hscHOs exhibited active spontaneous firing, which was significantly suppressed upon the application of 5-HT (Fig. 3c and Extended Data Fig. 7a). Moreover, pharmacological interventions with two selective 5-HT reuptake inhibitors (SSRIs), fluoxetine and escitalopram, also significantly reduced the firing rate in hsrHOs and hscHOs, suggesting that SNs in organoids are functionally responsive to SSRI-mediated blockade of 5-HT reuptake (Fig. 3c and Extended Data Fig. 7b, c). In addition, treatment with muscimol, a GABA_A_ receptor agonist, and L-glutamic acid, the principal excitatory neurotransmitter, resulted in corresponding decreases or increases in the mean firing rate, respectively, suggesting the presence of functional GABAergic and glutamatergic neurotransmission within the organoids (Extended Data Fig. 7d, e). Furthermore, immunostaining and quantification confirmed a significant increase in the number of c-FOS^+^ cells in hsrHOs and hscHOs following KCl-induced depolarization, indicating robust neural excitability and activity-dependent gene expression (Fig. 3d).

**Fig. 3.**
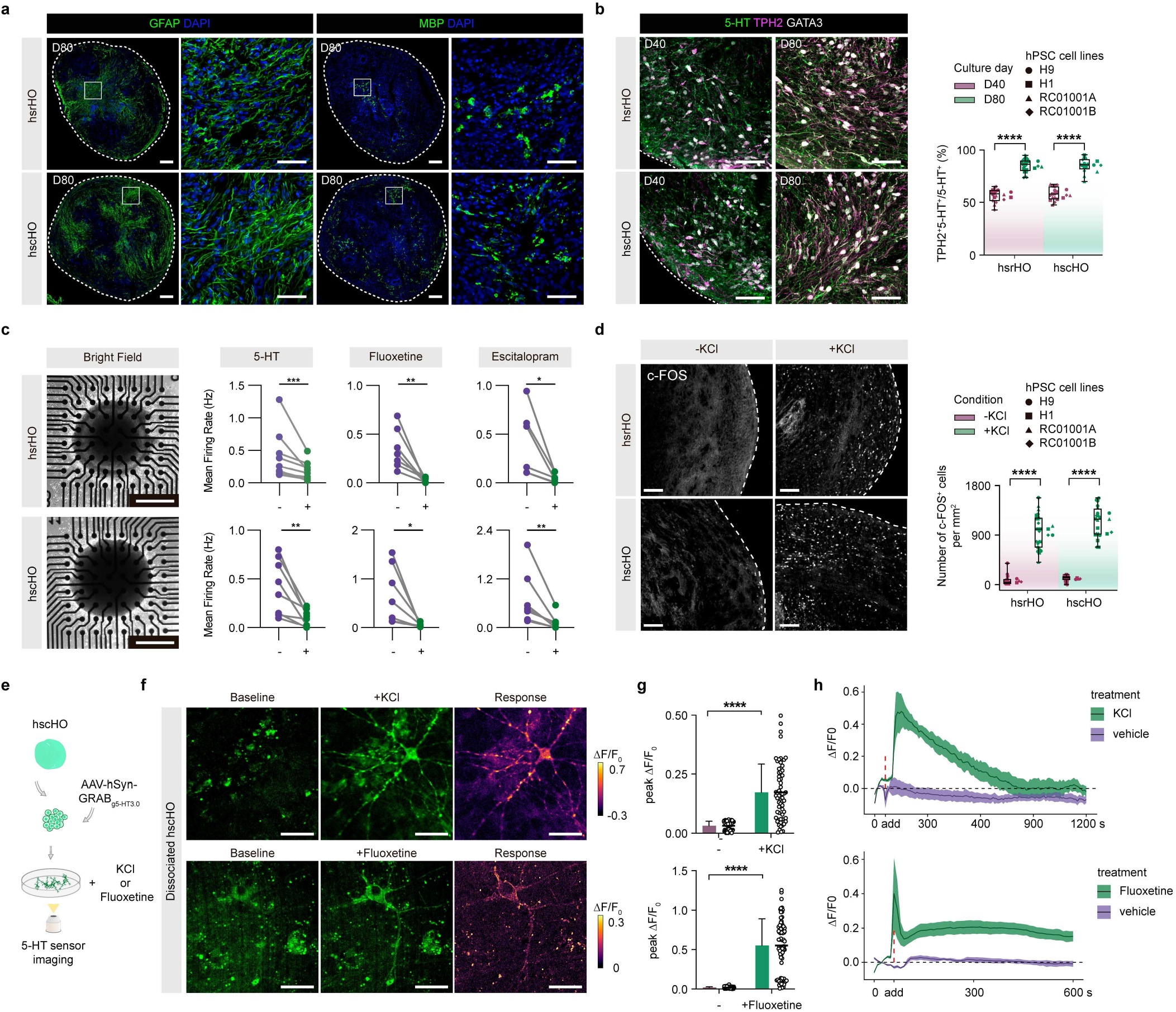
Functional development of hsrHOs and hscHOs. **a**, Representative immunostaining images of astrocytes (GFAP^+^) and oligodendrocytes (MBP^+^) at day 80 of ALI-cultured hsrHOs and hscHOs. **b**, Representative immunostaining images and quantification of mature SNs (5-HT^+^TPH2^+^ double-positive) at day 40 and day 80 in hsrHOs and hscHOs (hsrHOs, n = 14 of day 40 and n = 21 of day 80 derived from 4 hPSC cell lines, two to four differentiations, ****p < 0.0001, two-tailed Welch’s t test; hscHOs, n = 15 of day 40 and n = 13 of day 80 derived from 4 hPSC cell lines, two to four differentiations, ****p < 0.0001, two-tailed Welch’s t test). **c**, MEA assays of hsrHO and hscHO sections under spontaneous conditions and treatment of 5-HT, fluoxetine, or escitalopram (5-HT, n = 9 of hsrHOs, ***p = 0.0003, two-tailed ratio paired t test, n = 8 of hscHOs, **p = 0.0072, two-tailed paired t test; Fluoxetine, n = 7 of hsrHOs, **p = 0.0083, two-tailed ratio paired t test, n = 7 of hscHOs, *p = 0.0270, two-tailed paired t test; Escitalopram, n = 6 of hsrHOs, *p = 0.0331, two-tailed paired t test, n = 7 of hscHOs, **p = 0.0024, two-tailed ratio paired t test). **d**, Representative immunostaining images and quantification of c-FOS-positive cells in hsrHOs and hscHOs following KCl-induced depolarization (hsrHOs, n = 23 of before and n = 23 of KCl treatment from 4 hPSC cell lines, two to four differentiations, ****p < 0.0001, two-tailed lognormal Welch’s t test; hsrHOs, n = 29 of before and n = 20 of KCl treatment from 4 hPSC cell lines, two to four differentiations, ****p < 0.0001, two-tailed Welch’s t test). **e**, Schematic diagram of the 5-HT sensor imaging experiment. **f**, Imaging of 5-HT sensor fluorescence intensity changes under spontaneous conditions and upon treatment with KCl or fluoxetine in dissociated hscHOs expressing AAV9-hSyn-GRAB_g5-HT3.0_. **g**, Quantification of 5-HT sensor fluorescence intensity changes for spontaneous acitivity and treatment with KCl or fluoxetine (KCl, n = 62 cells from six independent experiments, ****p < 0.0001, two-tailed Mann-Whitney test; Fluoxetine, n = 75 cells from six independent experiments, ****p < 0.0001, two-tailed Mann-Whitney test). **h**, Representative time-lapse recording of 5-HT sensor fluorescence intensity changes under spontaneous conditions and upon KCl or fluoxetine treatment (KCl, n = 4 cells for KCl and n = 4 for vehicle; Fluoxetine, n = 12 for fluoxetine and n = 7 for vehicle). Data were presented as mean ± SD. Scale bars, 200 μm (a), 50 μm (a(zoom), b, f), 500 μm (c), 100 μm(d).

To evaluate the 5-HT release capacity of SNs, day-80 hscHOs were dissociated into single cells, reseeded, and transduced with pAAV9-hSyn-GRAB_g5-HT3.0_^39^ to express the GPCR-activation-based 5-HT sensor for monitoring 5-HT levels (Fig. 3e). Upon neuronal depolarization induced by KCl or treatment with SSRI fluoxetine, the fluorescence intensity of GRAB_g5-HT3.0_ increased significantly, indicating that SNs are capable of releasing 5-HT (Fig. 3f, g). Besides, continuous recordings showed that fluorescence in the KCl-treated group peaked and returned to baseline within 15 min, whereas in the fluoxetine-treated group exhibited a sustained elevation in fluorescence over time (Fig. 3h), consistent with the transient depolarizing effect of KCl and the sustained blockade of 5-HT reuptake by fluoxetine. These results demonstrate that hsrHOs and hscHOs acquire functional serotonergic neuronal properties during long-term culture.

### Distinct projection preferences of SNs in hsrHOs and hscHOs

Recent studies have shown that SNs in different segments of the hindbrain possess distinct projection targets throughout the nervous system^9,40,41^. Specifically, rostral SNs project broadly to the forebrain and midbrain, whereas caudal SNs preferentially extend descending projections to the spinal cord. Consistent with their identity, our scRNA-seq data revealed that SNs in hscHOs were predominantly TAC1-positive, a marker for a specific subgroup known to project axons to the ventral spinal cord, rather than EGR2-positive, a marker for a specific subgroup known to project axons to the dorsal spinal cord^2^ (Extended Data Fig. 8a). To understand whether *in vitro* organoid-derived SNs remain their projection preferences, we transduced hsrHOs and hscHOs with AAV9-TPH2::mCherry vectors and fused them with hCOs and human ventral spinal cord organoids (hvSpOs) (Fig. 4a and Extended Data Fig. 8b-d). 30 days post fusion (dpf), these two types of assembloids (hCO-hsrHO-hvSpO and hCO-hscHO-hvSpO) were imaged to quantify serotonergic axonal projections. Notably, we found that TPH2::mCherry^+^ serotonergic axons from hsrHOs preferentially extended toward hCOs rather than hvSpOs, whereas those from hscHOs showed the opposite preference, projecting more robustly toward hvSpOs than hCOs; the distinct axonal projection preferences of hsrHOs and hscHOs were reproducibly observed in assembloids generated from multiple cell lines (Fig. 4b, c and Extended Data Fig. 8e, f). Quantification across four hPSCs lines further demonstrated these differences in projection density (Fig. 4d), indicating that *in vitro* differentiated human hindbrain segment-specific SNs preserve *in vivo* like projection behaviors.

**Fig. 4.**
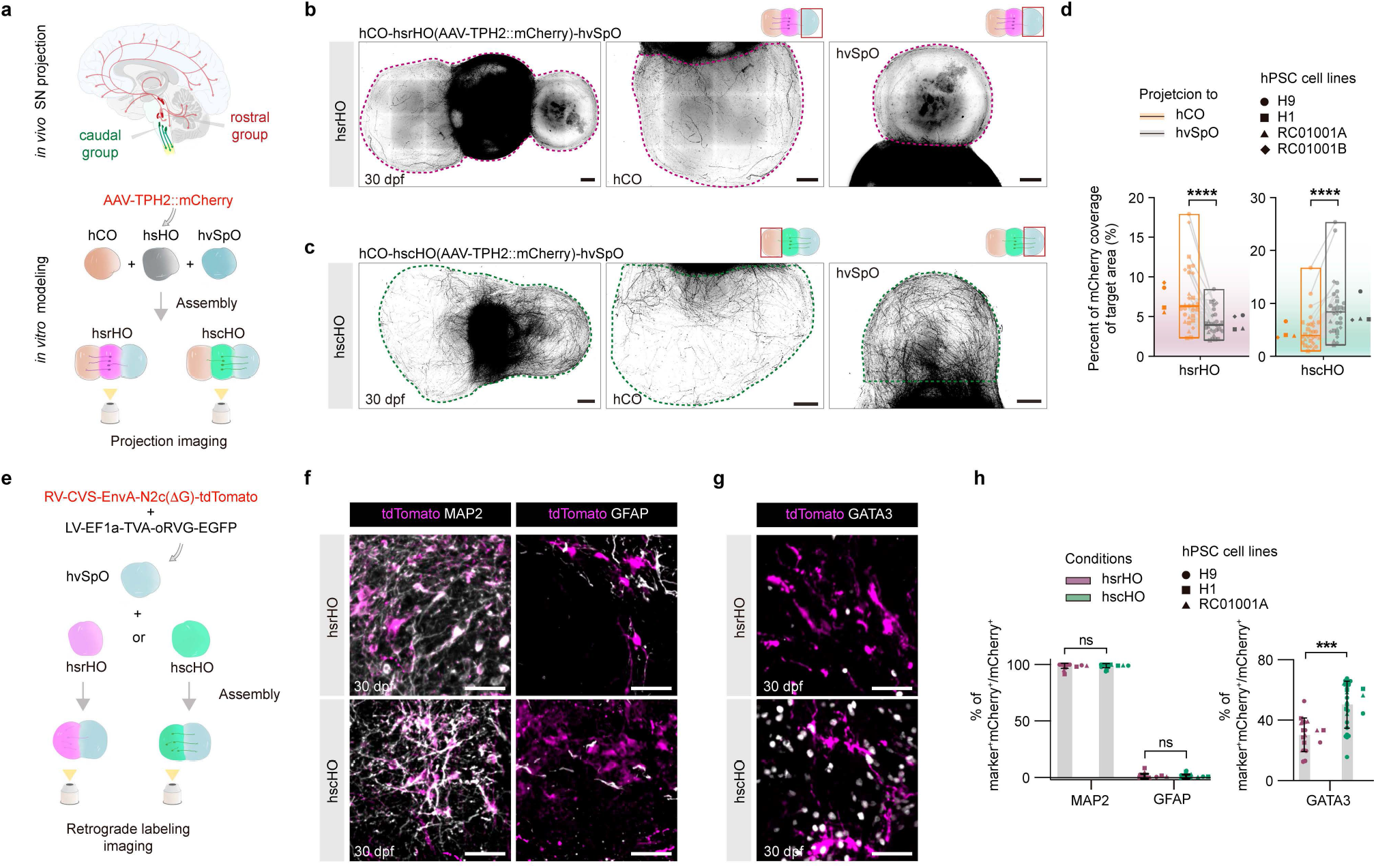
Projection patterns of SNs in hsrHOs and hscHOs. **a**, Schematic diagram of SN projection patterns *in vivo* (top) and *in vitro* modeling using hCO-hsrHO-hvSpO and hCO-hscHO-hvSpO assembloids (bottom). **b-d**, Representative images of serotonergic axon projections (mCherry^+^) from hsrHO (**b**) or hscHO (**c**) to hCO and hvSpO, and quantification of mCherry fluorescence coverage of hCO and hvSpO (**d**) (hCO-hsrHO-hvSpO, n = 29 assembloids from four hPSC cell lines, three to five differentiations, ****p < 0.0001, two-tailed ratio paired t test; hCO-hscHO-hvSpO, n = 31 assembloids from four hPSC cell lines, three to five differentiations, ****p < 0.0001, two-tailed ratio paired t test). **e**, Schematic diagram of rabies-ΔG tracing experiment. **f**, Immunostaining of tdTomato with neuron marker MAP2 or astrocytes marker GFAP in hsrHOs or hscHOs. **g**, Immunostaining of tdTomato with SNs marker GATA3 in hsrHOs or hscHOs. **h**, Quantification of cells positive for tdTomato and MAP2, GFAP, or GATA3 in hsrHOs or hscHOs (MAP2, n = 17 hsrHO-hvSpO and n = 21 hscHO-hvSpO assembloids from three hPSC cell lines, two to three differentiations, p > 0.9999, two-tailed Mann-Whitney test; GFAP, n = 17 hsrHO-hvSpO and n = 21hscHO-hvSpO assembloids from three hPSC cell lines, two to three differentiations, p > 0.9999, two-tailed Mann-Whitney test; GATA3, n = 16 hsrHO-hvSpO and n = 24 hscHO-hvSpO assembloids from three hPSC cell lines, two to three differentiations, ***p = 0.0003, two-tailed Mann-Whitney test). Scale bars, 500 μm (b, c), 50 μm (f, g).

To further investigate the projection behavior of SNs, we focused on descending projections and performed retrograde tracing experiments (Fig. 4e). hsrHOs and hscHOs were fused with hvSpOs that had been pretransduced with RV-CVS-EnvA-N2c(ΔG)-tdTomato and LV-EF1a-TVA-P2A-oRVG-P2A-EGFP-CMV-Puro-WPRE, and imaging analysis was performed at 30 dpf. We found that retrogradely traced tdTomato^+^ cells in hsrHOs and hscHOs were predominantly MAP2^+^ neurons and only rarely GFAP^+^ glial cells (Fig. 4f, h). Consistent with the distinct projection preferences between hsrHOs and hscHOs (Fig. 4b-d and Extended Data Fig. 8e, f), hscHOs contained a significantly higher portion of retrogradely traced GATA3^+^tdTomato^+^ SNs than hsrHOs, indicating that caudal SNs connect with cells in hvSpOs and preferentially project toward spinal cord region compared with hsrHOs (Fig. 4g, h). Together, our results demonstrate that hsrHOs and hscHOs containing segment-specific SNs recapitulate *in vivo*-like target-specific axonal projection preferences.

### Serotonergic motor control in human hindbrain-spinal-muscular assembloid

By providing an essential neuromodulatory drive that gates and tunes spinal motoneuron excitability, hindbrain SNs critically support coordinated locomotor activity^1,42^; notably, this modulation follows a non-monotonic pattern, where moderate 5-HT facilitates movement while excessive release promotes central fatigue, a state where the central nervous system fails to maintain the necessary drive to the muscles^1,43,44^. Modeling serotonergic control of motor function in human systems has been challenging due to the lack of established human models. To establish the human serotonin–spinal cord–muscle axis *in vitro* and investigate how hindbrain-derived 5-HT influences muscle control, we generated human hindbrain–spinal–muscular assembloids (hHSMAs) by fusing hscHOs with our previously established neuromusculoskeletal tri-tissue organoids (hNMSOs), in which the R1 neural region possesses a ventral spinal cord identity and forms functional neuromuscular junctions with the R2 muscle region^25^ (Fig. 5a, b). We utilized hscHOs because, as observed both *in vivo*^4,40,41^ and in our *in vitro* model (Fig. 4b-d and Extended Data Fig. 8e, f), SNs originating from the caudal hindbrain are the primary source of descending serotonergic projections to the spinal cord. The structure of both hscHO and hNMSO were preserved in hHSMAs and serotonergic axons from hscHO projected extensively to the R1 region of hNMSO (Fig. 5b, c). Notably, we found that assembling hscHOs with the neural domain of hNMSOs significantly increased the median amplitude of calcium spikes in skeletal muscle cells (SMCs) in the hNMSO R2 muscle region (Fig. 5d, e), suggesting that integration with hindbrain components promotes muscle function. To validate the contribution of serotonergic signaling to these effects, we applied the 5-HT2A receptor antagonist cyproheptadine (CPH), which significantly attenuated the observed enhancement (Fig. 5d, e). In contrast, in hNMSOs that were not fused with hscHOs, skeletal muscle cell activity remained lower than in hHSMAs and showed no significant change upon CPH treatment (Fig. 5d, e). These results indicate that 5-HT released from hscHOs enhances skeletal muscle functionality.

**Fig. 5.**
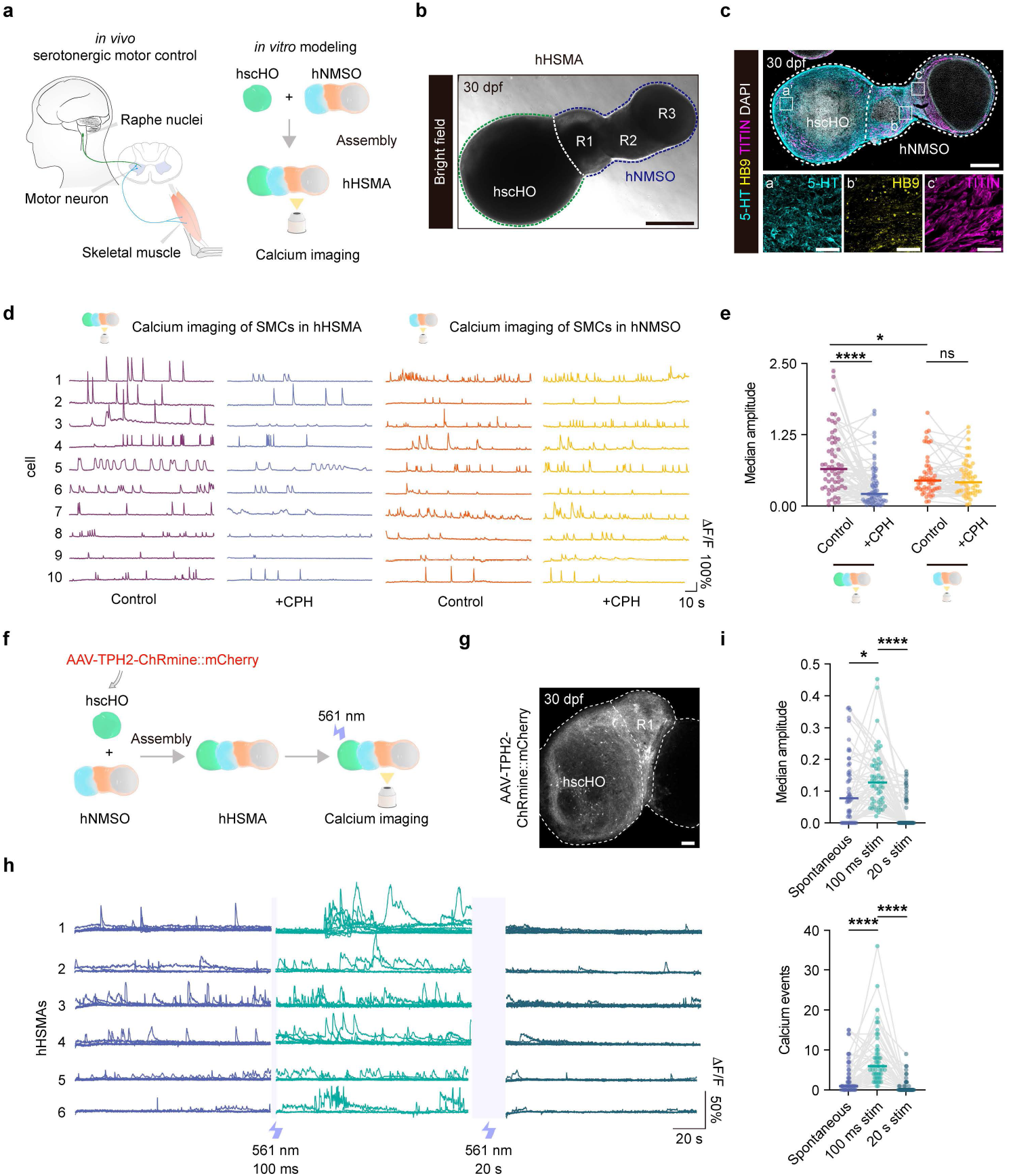
Serotonergic motor control in hHSMAs. **a**, Schematic diagram of 5-HT released by SNs modulating motor outputs *in vivo* (top) and *in vitro* modeling using hscHO-hNMSO assembloids (hHSMAs) (bottom). **b-c**, Bright field (**b**) and immunostaining (**c**) images of hHSMAs for 5-HT, HB9 and TITIN. **d-e**, Calcium imaging of spontaneous and cyproheptadine (CPH)-treated activities in skeletal muscle cells of hHSMAs or hNMSOs (**d**), and quantification (**e**) (for SMCs in hHSMAs, n = 61 cells from 10 hHSMAs of 3 batches, ****p < 0.0001, two-tailed Wilcoxon test; for SMCs in hNMSOs, n = 51 cells from 12 hNMSOs of 2 batches, p = 0.3314, two-tailed Wilcoxon test; for cells in hHSMAs and hNMSOs before CPH treatment, *p = 0.0181, two-tailed Mann-Whitney test). **f**, Schematic diagram of calcium imaging of SMCs in hHSMAs under different photostimulation conditions. **g**, Imaging of AAV9-TPH2-ChRmine::mCherry expression in hscHOs and their axon projections (mCherry^+^) to hNMSOs in hHSMAs. **h-i**, Calcium imaging of spontaneous activities and responses to 100 ms or 20 s optogenetic stimulation in hHSMAs (**h**), and quantification (**i**) (n = 53 cells from 13 hHSMAs of four batches; median amplitude, *p = 0.0175 for spontaneous versus 100 ms stimulation and ****p < 0.0001 for 20 s stimulation versus 100 ms stimulation, one-way ANOVA, Friedman test for multiple comparisons; calcium events, **** p < 0.0001 for spontaneous versus 100 ms stimulation and ****p < 0.0001 for 20 s stimulation versus 100 ms stimulation, one-way ANOVA, Friedman test for multiple comparisons). Scale bars, 500 μm (b, c), 200 μm (g), 50 μm (c(zoom)).

To further evaluate serotonergic control of motor activity in hHSMAs, we transduced hscHOs with AAV9-TPH2-ChRmine::mCherry to drive the expression of opsin ChRmine specifically in SNs. These were then assembled with hNMSOs and subjected to functional imaging at 30 dpf (Fig. 5f, g). To assess both the acute and prolonged effects of hscHO stimulation, we recorded muscle calcium transients throughout a continuous session comprising a brief 100 ms optogenetic pulse followed by 20 s of sustained stimulation specifically targeting the hscHO SNs. We found that while SMCs displayed spontaneous calcium activity, the 100 ms pulse elicited a significant increase in calcium spike amplitude; conversely, following the 20 s photostimulation, these calcium surges were nearly abolished (Fig. 5h, i), suggesting that hHSMAs can recapitulate the 5-HT-mediated transition from facilitatory modulation to central fatigue^1,45^. Collectively, these findings demonstrate that the serotonergic modulation of motor function can be modeled within the hHSMA system.

### DISC1 deficiency impairs the human serotonergic–motor axis

We next investigated whether our model could be utilized to recapitulate human serotonergic dysfunction. To this end, we focused on schizophrenia (SCZ), a severe neurodevelopmental disorder associated with serotonergic abnormalities^46-48^. Despite this potential link, the specific effects of the disorder on the human hindbrain serotonergic system remain poorly understood, primarily due to a lack of relevant human-specific models. To model SCZ, we generated DISC1-knockout hESCs using CRISPR-mediated gene editing, targeting DISC1, a major schizophrenia risk gene^49,50^. Ultimately, we established two wild-type (WT) cell lines and two knockout (KO) cell lines (Fig. 6a and Extended Data Fig. 9a). Both WT and DISC1 KO hESC lines were subsequently differentiated into hscHOs. Across multiple developmental stages, from day 16 to day 90, DISC1 KO hscHOs were significantly reduced in size compared with WT controls (Extended Data Fig. 9a, b). Compared with WT hscHOs, DISC1 KO hscHOs showed a significant reduction in the proportions of GATA3^+^ SNs, and the remaining neurons exhibited persistent abnormal somatic hypertrophy throughout development from day 40 to day 90 (Fig. 6b, c), indicating that DISC1 deficiency impairs the differentiation of human hindbrain SNs. To investigate potential alterations in the human serotonergic–motor axis, we generated hHSMAs by fusion hscHOs from WT and DISC1-KO cell lines with hNMSOs. All hNMSOs used for assembling hHSMAs were derived from un-edited hESCs to exclude potential effects of the DISC1 mutation on muscle development. Before assembly, hscHOs were transduced with AAV9-TPH2::mCherry vectors (Fig. 6d). At 30 dpf, quantification of mCherry+ serotonergic projections in the ventral spinal cord region of hNMSOs (R1) showed a significant reduction in projections derived from DISC1 KO hscHOs compared with WT controls (Fig. 6e). Functionally, consistent with the atypical serotonergic innervation pattern, we observed a significantly lower median amplitude of calcium transients within the SMCs of DISC1 KO hHSMAs compared with WT controls (Fig. 6f). These findings indicate that while SNs are modulators of motor output (Fig. 5a-e), DISC1 deficiency leads to impairments of this regulatory axis.

**Fig. 6.**
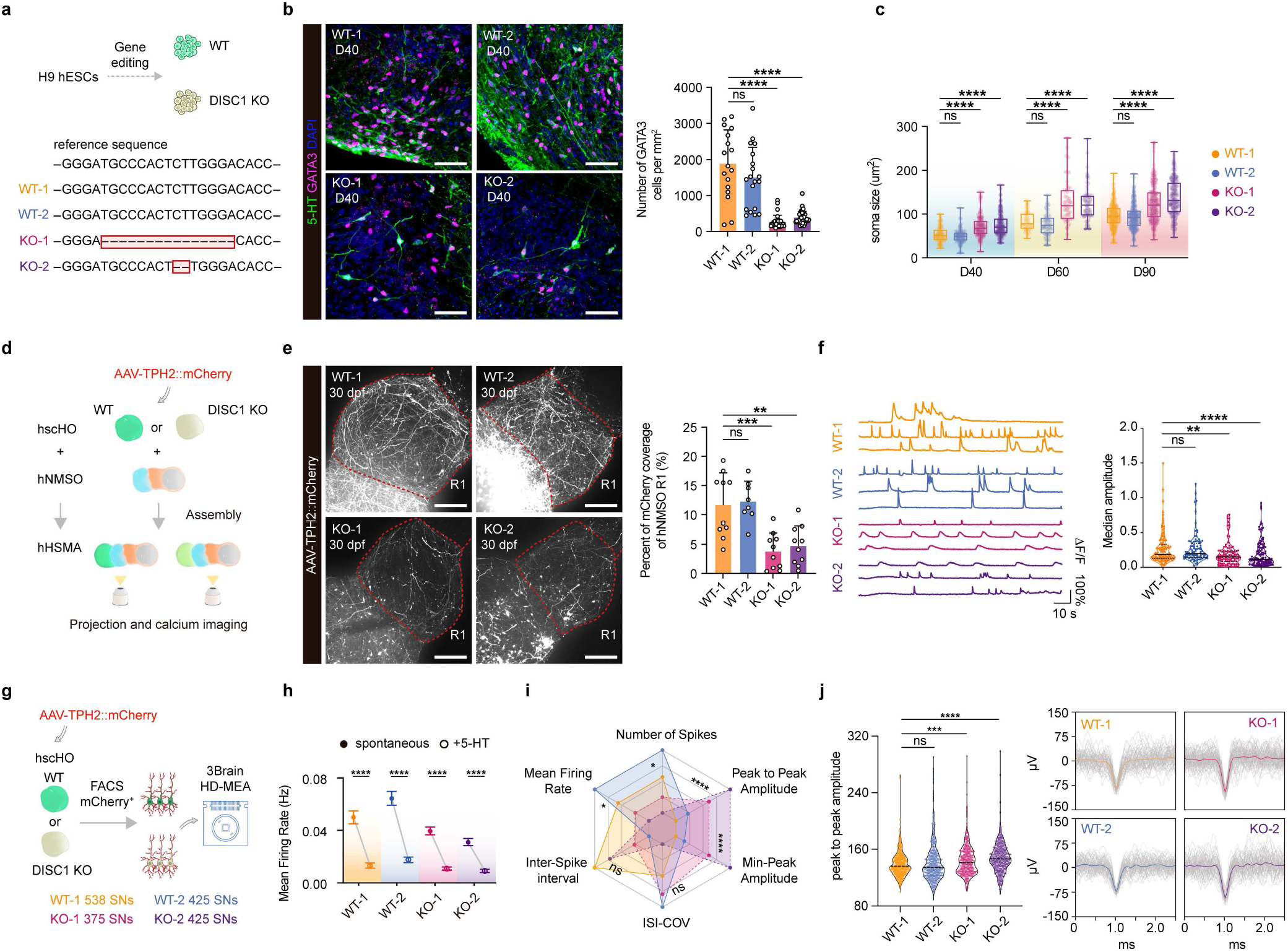
DISC1 deficiency impairs the human serotonergic–motor axis. **a**, Sanger sequencing of exon 2 of DISC1 gene in WT and DISC1 KO hPSC lines. **b**, Immunostaining and quantification of 5-HT^+^GATA3^+^ positive SNs in day 40 hscHOs generated from WT and DISC1 KO hPSC lines (WT-1, n = 17 from three differentiations; WT-2, n = 19 from three differentiations; KO-1, n = 29 from five differentiations; KO-2, n = 37 from five differentiations; p > 0.9999 for WT-2 versus WT-1, ****p < 0.0001 for KO-1 versus WT-1, ****p < 0.0001 for KO-2 versus WT-1, one-way ANOVA, Kruskal-Wallis test for multiple comparisons). **c**, Quantification of SN soma size in hscHOs generated from WT and DISC1 KO hPSC lines at day 40, 60, and 90 (For day 40: WT-1, n = 100; WT-2, n = 109; KO-1, n = 136; KO-2, n = 177; three to five differentiations, p = 0.3791 for WT-2 versus WT-1, ****p < 0.0001 for KO-1 versus WT-1, ****p < 0.0001 for KO-2 versus WT-1, one-way ANOVA, Kruskal-Wallis test for multiple comparisons. For day 60: WT-1, n = 45; WT-2, n = 69; KO-1, n = 50; KO-2, n = 66; two to four differentiations, p = 0.7746 for WT-2 versus WT-1, ****p < 0.0001 for KO-1 versus WT-1, ****p < 0.0001 for KO-2 versus WT-1, one-way ANOVA, Kruskal-Wallis test for multiple comparisons. For day 90: WT-1, n = 245; WT-2, n = 241; KO-1, n = 197; KO-2, n = 163; three to five differentiations, p = 0.7030 for WT-2 versus WT-1, ****p < 0.0001 for KO-1 versus WT-1, ****p < 0.0001 for KO-2 versus WT-1, one-way ANOVA, Kruskal-Wallis test for multiple comparisons). **d**, Schematic diagram describing the generation of hHSMAs using hNMSOs derived from H9 cell line and hscHOs derived from WT and DISC1 KO hPSC lines and infected with AAV9-TPH2::mCherry vectors. **e**, Imaging and quantification of axon projections (mCherry^+^) from hscHOs to R1 (ventral spinal cord) region in hHSMAs (WT-1, n = 10 hHSMAs; WT-2, n = 8 hHSMAs; KO-1, n = 10 hHSMAs; KO-2, n = 10 hHSMAs; two to four differentiations, p = 0.9823 for WT-2 versus WT-1, ***p = 0.0003 for KO-1 versus WT-1, **p = 0.0012 for KO-2 versus WT-1, ordinary one-way ANOVA). **f**, Calcium imaging of spontaneous activities for skeletal muscle cells in WT and DISC1 KO hHSMAs, and quantification (WT-1, n = 141 cells from 10 assembloids; WT-2, n = 91 cells from 7 assembloids; KO-1, n = 105 cells from 10 assembloids; KO-2, n = 125 cells from 9 assembloids; three to four differentiations, p > 0.9999 for WT-2 versus WT-1, **p = 0.0015 for KO-1 versus WT-1, ****p < 0.0001 for KO-2 versus WT-1, one-way ANOVA, Kruskal-Wallis test for multiple comparisons). **g**, Schematic diagram describing extracellular recording of WT and DISC1 KO SNs using the 3Brain HD-MEA system. **h**, Mean firing rate of WT and DISC1 KO SNs under spontaneous conditions and upon 5-HT treatment. Data are shown as mean ± SEM. (WT-1, n = 538 SNs from three differentiations, ****p < 0.0001; WT-2, n = 425 SNs from three differentiations; ****p < 0.0001; KO-1, n = 375 SNs from three differentiations, ****p < 0.0001; KO-2, n = 425 SNs from two differentiations, ****p < 0.0001; two-tailed Mann-Whitney test). **i**, Radar plot of spike features of WT and DISC1 KO SNs (number of spikes, *p = 0.0402 for WT versus KO; mean firing rate, *p = 0.0339 for WT versus KO; inter-spike interval, p = 0.1891 for WT versus KO; ISI-COV, p = 0.1016 for WT versus KO; min-peak amplitude, ****p < 0.0001 for WT versus KO; peak-to-peak amplitude, ****p < 0.0001 for WT versus KO). **j**, Violin plot of peak-to-peak amplitude and representative waveform of WT and DISC1 KO SNs (p > 0.9999 for WT-2 versus WT-1, ***p = 0.0009 for KO-1 versus WT-1, ****p < 0.0001 for KO-2 versus WT-1, one-way ANOVA, Kruskal-Wallis test for multiple comparisons). Scale bars, 50 μm (b), 200 μm (e).

To directly assess the electrophysiological properties of SNs generated in WT and DISC1-KO hscHOs, we performed high-density microelectrode array (HD-MEA) recordings (Fig. 6g). hscHOs were transduced with AAV9-TPH2::mCherry vectors at day 40. Subsequently, at approximately day 90, hscHOs were dissociated, and mCherry^+^ neurons were isolated via fluorescence-activated cell sorting (FACS) before being plated onto HD-MEA chips. Following a 5-minutes recording of spontaneous activity, we applied 5-HT to the cultures; only electrodes units that exhibited a reduction in mean firing rate were included in the subsequent analysis (Fig. 6h). This criterion was used to functionally identify mature SNs, as their activity is typically inhibited by 5-HT treatment^4^. Radar plot of single-neuron spike features demonstrated that DISC1 deficiency resulted in electrophysiologically distinct neuronal populations (Fig. 6i). DISC1 KO SNs exhibited impaired activity compared to WT SNs, as evidenced by significant reductions in both the number of spikes and the mean firing rate (Fig. 6i). Besides, DISC1 KO SNs exhibited significantly higher peak-to-peak amplitudes than WT SNs (Fig. 6j), a finding that aligns with their increased soma size (Fig. 6c), as extracellular spike amplitude is often influenced by neuronal morphology^51,52^. Together, these results reveal that DISC1 deficiency impairs human serotonergic modulation of motor control and indicate that hHSMAs provide an *in vitro* platform for studying diseases associated with the serotonergic–motor axis.

## Discussion

Neuromodulators, such as 5-HT, govern a diverse array of physiological functions by modulating neural circuit activity^10^. Within the central nervous system, the serotonergic system is a heterogeneous assembly of distinct 5-HT-producing neurons distributed along the rostrocaudal axis of hindbrain that project extensively to the brain and spinal cord. Despite its critical role in maintaining basic physiological homeostasis and its association with debilitating psychiatric disorders when dysfunctional, modeling the serotonergic system *in vitro* with human cells remains a significant challenge. Here, by precisely controlling rostrocaudal and ventral patterning, we developed differentiation methods to generate human hindbrain organoids from hPSCs with distinct r2–r3 rostral and r5–r8 caudal identities, respectively. These organoids successfully produced SNs that retained their respective regional identities (r2– r3 rostral or r5–r8 caudal). Our data showed that SNs in hsrHOs and hscHOs closely resemble *in vivo* SNs, offering a platform to study the development of human hindbrain segment-specific SNs. The generated SNs are capable of releasing 5-HT, as verified by a 5-HT sensor, and exhibited distinct response profiles to drug treatment, suggesting that organoid-derived SNs can serve as a platform for evaluating pharmacological responses. Notably, beyond their functional characteristics, we found that serotonergic projection preferences of rostral (hsrHOs) and caudal (hscHOs) SNs closely mirror those *in vivo*, thereby providing a human-relevant model for investigating the mechanisms underlying SN projection preferences, which has otherwise been challenging to study in a human system.

By fusing hscHOs with hNMSOs, we developed hHMSAs, an assembloid system that incorporates the key components of the human serotonergic–motor axis: SNs, motor neurons, and SMCs. To our knowledge, this represent the first *in vitro* model of human serotonergic motor control. This assembloid system thus provides an opportunity to dissect how hindbrain-derived serotonergic modulatory pathways regulate motor circuitry and downstream functional outputs in both physiological and pathological contexts. For instance, this model enables the investigation of how DISC1 deficiency impacts serotonergic motor control in human. This was also previously challenging to study in schizophrenia due to the lack of relevant models. In addition, given that serotonergic neurons have been shown to play a vital role in spinal cord injury repair^42,53-56^, these assembloid models, established by fusing hscHOs with spinal cord organoids or hNMSOs, could be leveraged not only to study serotonergic motor control dysfunction in disease, but also to investigate the mechanisms underlying spinal cord injury repair.

While hindbrain r2-r3 and r5-r8 regions give rise to distinct rostral and caudal SN populations, SNs derived from r1 region was not investigated in this study. The differentiation protocols for hsrHOs and hscHOs potentially can be extended to generate r1-derived SNs. Notably, a recent preprint by Kanton et al., described a human midbrain-hindbrain organoid (hMHO) model containing SNs that may recapitulate more rostral SNs identities than this study^57^. Together with our hsrHO and hscHO models, which represent the r2–r3 and r5–r8 identity, respectively, these systems collectively encompass much of the regional diversity of serotonergic neuronal subtypes in the brain.

Overall, we present a platform for generating human hindbrain organoids with segment-specific SNs and for modeling serotonergic modulatory pathways along the motor function axis. These models provide valuable tools for dissecting the development of human SNs and investigating how they establish and regulate serotonergic modulatory systems within target neural circuits.

## Methods

### Cell culture

Human ESC lines (H9 and H1) and iPSC lines (RC01001A and RC01001B) were maintained with mTeSR Plus media (Stem Cell Technologies, 100-0276) on growth factor-reduced Matrigel (Corning, 354230) coated cell culture plates. hESC lines H9 and H1 were commercially available from WiCell, and hiPSC lines RC01001A and RC01001B were commercially available from Nuwacell. RC01001A and RC01001B were derived via reprogramming of peripheral blood mononuclear cells of a 36-year-old male and umbilical cord cells of a newborn female, respectively. Cells were passaged every 4 or 5 days with 0.5 mM EDTA (Nuwacell, RP01007).

The use of hESCs and hiPSCs for all experiments in this work was approved by the ShanghaiTech Research Ethics Committee.

### Generation of hsrHOs and hscHOs

The protocol for generation of hsrHOs and hscHOs differ only during day 8 to day 12. Briefly, hPSCs were grown to a confluency of approximately 80% and dissociated with Accutase (Stem Cell Technologies, 07920). In total, 500 cells (for H9 cell line) were aggregated in ultra-low attachment 96-well U bottom plate (Corning, CLS7007-24EA) in hsHO induction media containing DMEM/F12 (Gibco, 11330057), 15% (v/v) KSR (Gibco, 10828028), 1% (v/v) MEM-NEAA (Gibco, 11140050), 1% (v/v) GlutaMAX (Gibco, 35050061) and 100 μM β-Mercaptoethanol (Sigma-Aldrich, M3148) supplemented with 10 μM SB431542 (Abcam, ab120163), 100 nM LDN (Sigma-Aldrich, SML0559), 3 μM CHIR99021 (Stem Cell Technologies, 72054) and 50 μM Y-27632 (MedChemExpress, HY-10583). From day 2 to day 8, change half of the hsHO induction media but not with Y-27632 every 2 days. At day 8, organoids were transferred to 6-well plates (BIOFIL, TCP030006) with 8 organoids/well to start spinning culture at 80 rpm/min on an orbital shaker (IKA, KS260). For hsrHOs, hsHO differenation media (1:1 mixture of DMEM/F12 and Neurobasal (Gibco, 21103049), 0.5% (v/v) N2 supplement (Thermo Fisher Scientific, 17502048), 1% (v/v) B27 supplement (Thermo Fisher Scientific, 17504044), 0.5% (v/v) MEM-NEAA, 1% (v/v) Glutamax, 1% (v/v) Penicillin/Streptomycin (Gibco, 15140122), 2.5 μg/mL Insulin (Beyotime, P3376), 50 μM β-Mercaptoethanol, 10 ng/mL EGF (Novoprotein, C029B), 20 ng/mL NGF (Novoprotein, C060)) supplemented with 100 ng/mL SHH (c25II, R&D, 464-SH-200) and 1 μM Purmorphamine (Stem Cell Technologies, 72204) was replenished every other day and 1% (v/v) dissolved Matrigel was added from day 8 to day 12. For hscHOs, the above-mentioned media supplemented with 100 nM RA (Sigma-Aldrich, R2625) was replenished every other day and 1% (v/v) dissolved Matrigel was added from day 8 to day 12. From day 12 to day 14, hsHO differenation media supplemented with 100 nM RA and 20 ng/mL FGF4 (Peprotech, 100-31-25UG) was used and replenished every other day. From day 16 to day 22, mature media (1:1 mixture of DMEM/F12 and Neurobasal, 0.5% (v/v) N2 supplement, 1% (v/v) B27 supplement, 0.5% (v/v) MEM-NEAA, 1% (v/v) Glutamax, 1% (v/v) Penicillin/Streptomycin, 2.5 μg/mL Insulin, 50 μM β-Mercaptoethanol, 20 ng/mL BDNF (Novoprotein, C076), 200 μM Ascorbic acid (Sigma-Aldrich, A92902)) supplemented with 2.5 μM DAPT (Abmole, M1746) and 2 ng/mL TGF-β2 (Abcam, ab84070) was used and replenished every other day. Beginning at day 22, mature media was replenished every other day before day 30 and every three days thereafter.

We used 500 cells per well for H9 hESCs line, 1000 cells per well for H1 hESCs line and 9000 cells per well for RC01001A and RC01001B hiPSCs lines for generation of hsHOs. For DISC1 WT and KO cell line, 1000 cells per well were used. To ensure reproducibility of generation in other cell lines, it is necessary to adjust the initial seeding cell number on the basis of the gene expression and immunofluorescence staining results.

### Generation of hCOs and hNMSOs

hCOs were generated as described previously with minor modifications^15^. Briefly, single-cell suspensions were plated in to ultra-low attachment 96-well U bottom plate (4500 cells per well for H9 hESCs) in the hCO induction media (DMEM/F-12, 15% KSR, 1% MEM-NEAA, 1% GlutaMAX and 100 μM β-Mercaptoethanol) supplemented with 10 μM SB431542, 100 nM LDN, 2 μM XAV939 (Sigma-Aldrich, X3004) and 50 μM Y-27632. From day 2 to day 10, change half of the hCO induction media but not with Y-27632 every other day. At day 10, organoids were transferred to 6-well plate with 8 organoids/well to start spinning culture. From day 10 to day 18, hCO differenation media (1:1 mixture of DMEM/F12 and Neurobasal, 0.5% (v/v) N2 supplement, 1% (v/v) B27 (-VA) supplement (Thermo Fisher Scientific, 12587010), 0.5% (v/v) MEM-NEAA, 1% (v/v) Glutamax, 1% (v/v) Penicillin/Streptomycin, 2.5 μg/mL Insulin, 50 μM β-Mercaptoethanol) was used and replenished every other day. On day 14 to day 18, 1% (v/v) dissolved Matrigel was added. Beginning at day 18, mature media was replenished every other day before day 30 and every three days thereafter.

hNMSOs were generated as described previously^25^. Briefly, single-cell suspensions were plated in to ultra-low attachment 96-well U bottom plate (100 cells/well for H9 line) in the mTeSR Plus media supplemented with 10 μM SB431542, 100 nM LDN, 3 μM CHIR99021 and 10 μM Y-27632. From day 2 to day 8, change half of the mTeSR Plus media supplemented with 100 nM RA and 20 ng/mL FGF2 but not with Y-27632 every other day. At day 8, organoids were transferred to 6-well plate with 8 organoids/well to start spinning culture. From day 8 to day 12, hNMSO differenation media (1:1 mixture of DMEM/F12 and Neurobasal, 0.5% (v/v) N2 supplement, 1% (v/v) B27 supplement, 0.5% (v/v) MEM-NEAA, 1% (v/v) Glutamax, 1% (v/v) Penicillin/Streptomycin, 2.5 μg/mL Insulin, 50 μM β-Mercaptoethanol) supplemented with 20 ng/mL FGF2, 100 nM RA and 100 nM SAG (Abcam, ab142160) was used and replenished every other day. From day 12 to day 16, hNMSO differenation media supplemented with 100 nM RA and 100 nM SAG was used and replenished every other day. From day 16 to day 24, mature media supplemented with 100 nM SAG was used and replenished every other day. Beginning at day 24, mature media was replenished every other day before day 30 and every three days thereafter.

### Immunofluorescence staining

Organoids were transferred to a 24 well-plate, washed 3 times with PBS and fixed in 4% PFA at 4°C overnight. Next, organoids were washed 3 times with PBS and incubated in 30% sucrose solution (in PBS) at 4°C for 2 days. Then, sucrose solution was removed and organoids were equilibrated in O.C.T compound (Sakura, 4583) at RT for 15 min. Organoids were transferred to a tissue base mold and embedded with O.C.T compound at -80°C and sectioned for 30 μm. The sections were incubated with 0.1% Triton-100 for 15 min and blocked with 3% bovine serum albumin (BSA) at RT for 2 hours and incubated with primary antibodies in 3% BSA at 4°C overnight. Sections were then washed 3 times with PBS and incubated with Alexa Fluor secondary antibodies (Invitrogen) diluted in 3% BSA at 1:1000 at RT for 1 hour. Sections were then washed 3 times with PBS and nuclei were stained with DAPI (Solarbio, C0065) diluted in PBS at 1:1000. Slides were mounted with CC/Mount (Sigma-Aldrich, C9368) and imaged with a Nikon CSU-W1 Sora 2Camera microscope and analyzed using Fiji (v2.16.0).

To assess neuron types and ratio in organoids, day 40 hsrHOs and hscHOs were dissociated with 0.25% Trypsin with EDTA (MeilunBio, MA0233-1), 0.1 mg/mL DNase I (MedChemExpress, HY-108882) at 37 °C for 15-20 min and gently mix every 5 min. Mature media was added to terminate digestion reaction and any undigested cell clumps were filtered out with a 70 μm cell strainer. Cells were resuspended with mature media supplemented with 10 μM Y-27632. The single-cell suspensions were reseeded on Matrigel coated confocal dishes and cultured about 2 weeks for immunofluorescence staining.

Primary antibodies used were: anti-NK2-2 (rabbit, 1:200; ab191077, Abcam), anti-NKX6-1 (rabbit, 1:200; ab221549, Abcam), anti-HOXA2 (rabbit, 1:200; NBP2-58865, Novus Biologicals), anti-HOXB4 (rabbit, 1:200; ab133521, Abcam), anti-SOX2 (rabbit, 1:500; 3579, Cell Signaling), anti-5-HT (goat, 1:10000; 20079, ImmunoStar), anti-TPH2 (rabbit, 1:500; NB100-74555, Novus Biologicals), anti-LMX1B (rabbit, 1:200; HPA073716, Atlas antibodies), anti-GATA3 (mouse, 1:500; MAB6330, R&D Systems), anti-GATA2/3 (rabbit, 1:500; ab182747, Abcam), anti-NR2F2 (mouse, 1:1000; ab41859; Abcam), anti-PAX7 (rabbit, 1:1000; ab92317, Abcam), anti-HOXC9 (mouse, 1:100; ab50839, Abcam), anti-FOXA2 (rabbit, 1:500; ab108422, Abcam), anti-NG2 (mouse, 1:2000; 554275, BD Biosciences), anti-GFAP (mouse, 1:500; MAB360, Millipore), anti-MBP (rat, 1:100; MAB386, Millipore), anti-c-FOS (rabbit, 1:100; 2250T, Cell Signaling), anti-mCherry (rabbit, 1:500; ab167453, Abcam), anti-mCherry (mouse, 1:500; ab125096, Abcam), anti-MAP2 (mouse, 1:1000; MAB3418, Millipore), anti-TH (rabbit, 1:1000; AB152, Millipore), anti-Tuj1 (mouse, 1:1000; ab78078, Abcam), anti-Tuj1 (rabbit, 1:1000; ab18207, Abcam), anti-vGlut2 (mouse, 1:500; MAB5504, Millipore), anti-GAD1 (mouse, 1:500; MAB5406, Millipore), anti-HB9 (mouse, 1:100, sc-515769, Santa Cruz), anti-TITIN (rabbit, 1:300, 27867, Proteintech).

### scRNA-seq and data processing

For single-cell RNA sequencing, day 40 hsrHOs and hscHOs were collected and pooled, respectively. Briefly, media was removed and organoids were washed with PBS. Then, organoids were cut with a surgical blade and dead cells inside the organoids were washed out with PBS. Organoids were dissociated into single cells in dissociation solution containing 0.25% Trypsin with EDTA, 0.1 mg/mL DNase I at 37 °C for 15-20 min and gently mix every 5 min. Add 1x PBS containing 2% (v/v) fetal bovine serum (FBS) to terminate digestion reaction. The cell suspension was then passed through a 70 μm cell strainer and centrifuged at 200g for 5 min. The cell viability should be above 80%. The single-cell suspensions were loaded to 10X Chromium to capture single cell according to the manufacturer’s protocol (v3.1 kit). For day 80 WT and DISC1 KO samples, GEM-X Single-Cell 3’ kit (v4) was used. The following cDNA amplification and library construction steps were performed according to the standard protocol. Libraries were sequenced on an Illumina NovaSeq 6000 platform (paired end, 150 bp) at a minimum depth of 20,000 reads per cell.

Cell Ranger (10x Genomic, v7.1.0) was used to map reads to human genome reference (GRCh38-2020-A). Next, the scRNA-seq data was analyzed using Seurat (v4.3.0) ^58^. Quality control was done by filtering cells with nFeature < 500 or > 7000, nCount < 2000 or > 25000 or mitochondrial transcript percentage higher than 15%. Doublets were then removed using DoubletFinder (v2.0.3) ^59^ with default parameter. Additional quality control step was removing stressed cell using Gruffi package (v1.5.5) ^60^.

For day 40 hsrHOs and hscHOs scRNA-seq data, gene expression was normalized using SCTransform function, top 2000 highly variable genes were detected and samples were integrated using Canonical Correlation Analysis (CCA) method implemented in Seurat. Principal Component Analysis (PCA) was performed and top 25 PC was used for dimensionality reduction in an UMAP embedding. Cell clusters were identified by shared nearest-neighbour graph construction and modularity detection using the FindNeighbors and FindClusters functions (dims = 30). We annotated cell classed through the expression of canonical markers and we further checked the markers of each cluster detected by FindAllMarkers function. Specifically, floor plate cells were identified by the expression of SHH, FOXA2 and SPON1; NPCs were identified by the expression of SOX2, NES and MIK67; OPCs were identified by the expression of CSPG4, SOX10 and PLP1, neural clusters were identified by the expression of INA and DCX. The neural clusters can be further divided into immature neurons (ASCL1^+^), GABAergic neurons (GAD1^+^GAD2^+^), glutamatergic neurons (SLC17A6^+^), visceral motor neurons (CHAT^+^TH^+^PHOX2B^+^) and serotonergic neurons (FEV^+^GATA3^+^LMX1B^+^). Markers of serotonergic neurons were identified by FindMarkers function compared to all other cell types using Wilcox test and Gene Ontology (GO) analysis was performed by clusterProfiler package (v4.9.0.2) ^61^. To construct the differentiation trajectory of cells in the hsHOs, we performed RNA velocity analysis using scvelo (v0.2.5) ^62^. To understand the SNs identity in the organoid, we mapped neural types to mouse E13.5 brain data through voxhunt package (v1.0.1) ^63^. We next integrated the organoid scRNA-seq data with serotonergic neuron transcriptome profiles extracted from human developing and adult brain atlas^64,65^. Specifically, we extract the FEV positive SNs from human developing stage data (https://github.com/linnarsson-lab/developing-human-brain/) and the cluster 397 which serotonergic markers FEV and SLC6A4 were highly expressed from the human adult stage data (https://github.com/linnarsson-lab/adult-human-brain) and manually annotated region information according to the original article. Then we used the anchor-based label transfer method to compare the transcriptome similarity between organoid and human data. The anchors were identified using FindTransferAnchors function with top 10 PCs and the prediction scores were summarized to each cell class as the similarity score. The differentially expression genes (DEGs) were identified using FindMarkers function and genes with absolute value of avg_log2FC > 0.5 and adjust p value < 0.01 were labeled as significantly regulated.

### Bulk RNA-seq library preparation and data processing

For bulk RNA-seq, day 40 hsrHOs and hscHOs were collected and pooled, respectively. Total RNA was extracted using the FastPure Cell/Tissue Total RNA Isolation Kit (Vazyme, RC101) according to the manufacturer’s protocol. RNA concentration and purity was determined using a nanodrop 2000 spectrophotometer and, RNA integrity was evaluated by Agilent Bioanalyzer 2100 with RIN number >7.0, and confirmed by electrophoresis with denaturing agarose gel. RNA samples passed quality control were used to constructed libraries using TruSeq Stranded mRNA LT Sample Prep Kit according to the manufacturer’s protocol. Lastly, 2×150bp paired-end sequencing (PE150) was performed on an illumina Novaseq™ 6000 platform following the manufacturer ‘s recommended protocol.

The nf-core/rnaseq pipeline^66^ (v3.22.2) was employed to quantitatively analyze gene expression. Briefly, clean reads were aligned to reference genome (GRCh38-3.0.0) using STAR^67^ (v2.7.10a) and RSEM^68^ (v1.3.1) was used to quantifying the expression of genes. Pearson correlation analysis was performed to assess the similarity between samples. The samples were normalized and DEGs were identified using DESeq2 package^69^. We used DWLS^70^ to deconvolute the cell type composition of bulk RNA-seq data using our day 40 scRNA-seq profile as reference.

### Real-time quantitative PCR

For RT-qPCR, at least 4 organoids were collected and pooled together. Total RNA was exacted as described above. cDNA was prepared by reverse transcription using the HiScript III All-in-one RT SuperMix Perfect for qPCR kit (Vazyme, R333) with 500 ng total RNA. qPCR was performed using Taq Pro Universal SYBR qPCR Master Mix (Vazyme, Q712) on QuantStudio 7 Real-Time PCR System (ThermoFisher Scientific) according to the manufacturer’s protocol. Primers used were listed in Extended Data Table 1.

### Air-liquid interface culture

For long-term culture of hsrHOs and hscHOs, we preformed vibratome sectioning on day 25-30 hsrHOs and hscHOs. Briefly, 4-6 organoids were collected and embedded in 3% low-gelling-temperature agarose (Sigma-Aldrich, A9414) in an embedding mold. Then the agarose block containing organoids were transfer to a vibrating microtome (Lecia, VT1200S) slot containing 1x PBS supplemented with 2% P/S. The organoids were sectioned into 300 μm thickness and slices were transferred to transwell plates (BIOFIL, TCS002006). Mature media was added to ensure sections remained at the air-liquid interface. The transwell plates with organoids sections were transferred onto an orbital shaker for long-term culture and mature media was change every other day.

### KCl treatment

For pharmacological treatment, KCl (Beyotime, ST345) was added to medium at a final concentration of 56 mM for 4 hours. Then organoids were collected and prepared for frozen section.

### 5-HT sensor GRAB_g5-HT3.0_ fluorescence imaging

To clear monitor changes in GRABg_5-HT3.0_ fluorescence intensity, organoids were dissociated into single cells as described above and plated onto confocal dishes (BIOFIL, BDD012035). On the second day, cells were infected with 300 nL of AAV9-hSyn-GRAB_g5-HT3.0_ (7.13 × 10^13^ vg/mL, WZ Biosciences, YL004015-AV9) per dish. After 2-3 weeks of recover and culture, living imaging was performed using a Nikon CSU-W1 Sora 2Camera microscope at 37 °C. Time-lapse images were acquired continuously at a rate of 10 second per frame for 10-20 min. The spontaneous fluorescence intensity was recorded for 1∼2 minutes before treatment. For pharmacological treatment, 56 mM KCl or 2 μM Fluoxetine (MedChemExpress, HY-B0102) was applied. Raw intensity was get using Fiji. Relative changes of fluorescence were calculated as ΔF/F_0_ = (F(t)-F_0_)/F_0_ where F_0_ was defined by the mean of spontaneous fluorescence intensity.

### Generation of hCO-hsrHO-hvSpO and hCO-hscHO-hvSpO assembloids and projection imaging

The sections of hCO, hsrHO, hscHO and hvSpO were used for generation of assembloids and sections of organoids were prepared as described above. Serotonergic neurons in around day 30 hsrHOs and hscHOs were labeled by infected with 150 μL AAV9-TPH2-mCherry (5.12 × 10^12^ vg/mL, BrainCase, BC-2179) per section. Then sections were placed onto transwell inserts and fused in the order of hCO-hsrHO-hvSpO and hCO-hscHO-hvSpO. After 30 days of fusion, the assembloid was transferred to a confocal dish and imaged using a Nikon CSU-W1 Sora 2Camera microscope at depth of 100-200 μm. Projection intensity was quantified in Fiji with the maximum-intensity merged images. To control for potential variables, the assembloids were generated using hsrHOs and hscHOs derived from different cell lines, whereas all hCOs and hvSpOs were generated from H9 cell line.

### Rabies-ΔG tracing

For rabies-ΔG tracing experiments, ∼ day 30 hvSpOs were transferred to ultra-low attachment 96-well U bottom plate with 150 μL fresh mature media. Both 60 nL of RV-CVS-EnvA-N2c(ΔG)-tdTomato (≥ 2.00 × 10^8^ IFU/mL, BrainVTA, R05002) and 200 nL of LV-EF1a-TVA-P2A-oRVG-P2A-EGFP-CMV-Puro-WPRE (≥ 5.00 × 10^8^ TU/mL, BrainVTA, LV-0478) were added. Two days after viral infection, hvSpOs were washed 3 times with fresh mature media. Then day 30 hsrHO or hscHO and hvSpO were assembled in one well of ultra-low attachment 96-well U bottom plate with 200 μL fresh mature media for 4 days. On day 2 of fusion, 100 μL mature media was gently replenished. Then assembloids were transferred to a 6-well plate onto an orbital shaker and mature media was change every other day until the assembloids were used for frozen section. Only sections with at least 10 tdTomato^+^ cells in the hsrHO or hscHO area were included in the analysis.

### Generation of hHSMAs

To generate hHSMAs, day 30 hscHOs and hNMSOs were used. Briefly, hscHO was placed onto transwell inserts to fusion with neural region of hNMSOs. After 2 days of fusion, the hHSMAs were carefully transferred with a Pasteur pipette into a 6-well plate and maintained on an orbital shaker. An ideal fused state was individual components of the hHSMA remained clearly distinguishable rather than appearing clumped together. The mature medium was changed every 2-3 days with maturation medium until the hHSMAs were used for experiments.

For photostimulation experiments, 300 nL of AAV9-TPH2-ChRmine::mCherry (3.10 × 10^12^ vg/mL, BrainCase, BC-1358) was injected into hscHO using RWD R480 glass microelectrode injection system. After 24 hours of injection, hscHOs were washed twice using fresh mature media and then fusion with neural region of hNMSOs as described above.

For disease modeling, to exclude potential effects of DISC1 mutation on muscular development, hHSMAs were generated by assembling hsrHOs derived from WT or DISC1 KO H9 hESCs with hNMSOs derived exclusively from un-edited H9 hESCs.

### Calcium imaging and photostimulation

Assembloids were transferred to a 12-well plate and incubated in BrainPhys Neuronal Medium (Stem Cell Technologies, 05790) containing 5 μM Cal-520 (AAT Bioquest, 21130) and 1% powerload (Thermo Fisher Scientific, P10020) at 37°C with 5% CO_2_ on an orbital shaker for 40 min. Then the media was replaced with fresh BrainPhys Neuronal Medium for another 20 min incubation. For recording, the assembloids were transferred to a confocal dish and calcium imaging was performed with a 4X or 10X objective at 488-nm excitation under a Nikon CSU-W1 Sora 2Camera microscope at 37°C. Time-lapse images were acquired continuously at a rate of 5 frames per second for 2 min. For drug treatment, 10 μM cyproheptadine (MedChemexpress, HY-B1622) was applied.

For the photostimulation experiments, we first defined an ROI over the TPH2-ChRmine::mCherry-labeled region of the hscHO and then performed continuous recording. We initially recorded spontaneous calcium activity for 2 min, followed by a 2 min recording immediately after a 100 ms stimulation in ROI with a 561 nm laser, and another 2 min recording after the 20 s photostimulation. All stimulation and imaging were performed within the same field of view.

Registering ROI, tracing of single-cell calcium and exporting raw fluorescent intensity was performed using Fiji. Relative changes of fluorescence were calculated as ΔF/F_0_ = (F(t)-F_0_)/F_0_ where F_0_ was the lower 5^th^ percentile value of the session. Calcium events were detected whenever the ΔF/F_0_ crossed a threshold of 5 median absolute deviations (MAD) and only those with a sharp rise and slower decay would be included in statistical analysis.

### DISC1 gene editing

To obtain WT and DISC1 KO hESCs, the sgRNA was designed using CRISPick (Broad institute, https://portals.broadinstitute.org/gppx/crispick/public). The selected sgRNA (5’-GAGCAGGGTGTCCCAAGAGT-3’) was synthesized and clone into pSpCas9(BB)-2A-Puro (PX459) construct. 2x 10^6^ of single cells dissociated from H9 hESCs were resuspended with 100 μL hPSC Nucleofection Buffer (Nuwacell, RP01005) and were electroporated with 5 μg of PX459-sgRNA using the electroporator (LONZA AAB-1001 Nucleofector™ 2BDevice) with B-016 program. The cells were then plated onto Matrigel-coated plate. Three days after electroporation, 1 μg/mL puromycin (Beyotime, STT551) was applied for 4 days and then cells were allowed for recovery for another 5 days. Then cells were dissociated into single cells and seeded in Matrigel-coated 96-well plate at 0.5 cell per well. After another 5∼7 days, single clones were picked manually and expanded. The sanger sequencing was used for validation and two WT and two DISC1 KO clones were picked for downstream experiments.

### Multielectrode Array (MEA) assay

Extracellular electrical recording of hsHO sections using the Axion Biosystems (M384-tMEA-6W) was performed as described before^25^. Briefly, around day 80 hsHO sections were plated on Matrigel coated electrode area in MEA plate and a drop of Matrigel was used to cover the section. After 30 min incubation at 37°C, 1 mL mature media was gently added to each well of MEA plate. The sections were cultured statically at 37°C with 5% CO2 and medium was changed every 2 days until recording after one week. Electrical recording was performed using Maestro pro MEA system and anaylzed with AxIS Navigator software (v3.7.1.16, https://www.axionbiosystems.com). Spontaneous activities were recorded for 2 min before 40 μM 5-HT (MedChemexpress, HY-B1473A), 2 μM fluoxetine, 5 μM escitalopram (MedChemexpress, HY-14258), 50 μM muscimol (MedChemexpress, HY-N2313) or 50 μM L-glutamic acid (MedChemexpress, HY-14608R) was applied. The mean firing rate was used for statistical test and Neural Metric Toolsoftware (v4.1.5, https://www.axionbiosystems.com) was used to plot electrode array activity.

For High-Density Microelectrode Array (HD-MEA) recording, ∼ day 40 organoids were infected with AAV9-TPH2-mCherry virus by injection 150 μL virus per organoid using RWD R480 glass microelectrode injection system. The organoids were then cultured in a 6 well dish on an orbital shaker to around day 90. Then organoids were dissociated as described above. The single cell suspension was then passed through a 70 μm cell strainer and centrifuged at 200g for 5 min. Then cells were resuspended with 3% BSA supplemented with 10 μM Y-27632 and put on ice for isolating mCherry^+^ cells by FACS using BD FACSAria III. After sorting, cells were centrifuged at 200g for 5 min and resuspended with mature media supplemented with 10 μM Y-27632 to 500 cells/μL. Then 60 μL cell suspension was seeded on Matrigel coated 2D MEA chip (CorePlate™ 1W) according to manufacturer’s protocol. After 4 hours of static culture at 37°C and 5% CO_2_, cells should be adhered to the chip and 500 μL mature media supplemented with 10 μM Y-27632 was then added. After 24 hours, 1 mL mature media was added. Cells were cultured for at least 2 weeks and the media was changed every 2 days until recording.

The recording was then performed using BioCAM DupleX (3Brain). Spontaneous activities were recorded for 5 min and then 40 μM 5-HT was applied to suppress the electrical activities of serotonergic neurons and recorded for another 5 min. Raw data was analyzed using BrainWave6 software (v6.0.9566.18925). To get high-quality electrical recording data, we used balanced mode for spike detection, filtered using mean firing rate ≥0.01 and electrical data with non-typical extracellular spike waveforms were manually filtered out. To make sure we are analyzing data of SNs, only data in which the mean firing rate after 5-HT application was lower than that during spontaneous recording were included in the analysis. Finally, we get high quality single-neuron resolution data after filtering (538 SNs from WT-1 hscHOs; 425 SNs from WT-2 hscHOs; 375 SNs from KO-1 hscHOs; 425 SNs from KO-2 hscHOs). Spike data including “Number of Spikes”, “Mean Firing Rate”, “Inter-Spike interval (ISI)”, “ISI -COV”, “Min Peak Amplitude (Abs value)”, “Peak-to-Peak Amplitude” was used for radar plot visualization.

### Statistics

Data were presented as mean ± SD unless otherwise stated. The paired or unpaired two-tail t-test and multiple comparison with one-way ANOVA or two-way ANOVA test were used to determine the statistical significance compare between groups. For data do not pass Shapiro-Wilk normality test, non-parametric test was used. GraphPad Prism (v11.0.1) was used for statistical analyses.

## Data availability

Gene expression data are available in the Gene Expression Omnibus (GEO) under accession numbers GSE330616. As of publication, data will be available without the need for a token.

## Acknowledgements

This study was supported by the National Key Research and Development Program of China (2024YFA1108000), the Joint Project of the Yangtze River Delta Science and Technology Innovation Community (2024CSJZN0600), the Lingang Laboratory (Grant No.LGL8998-11), the Central Guidance on Local Science and Technology Development Fund (YDZX20233100001002), the Shanghai Frontiers Science Center for Biomacromolecules and Precision Medicine at ShanghaiTech University, the Institute Development Fund of ShanghaiTech University, and the ShanghaiTech start-up fund. We thank the facility support from the Molecular Imaging Core and the Molecular and Cell Biology Core at the School of Life Science and Technology, ShanghaiTech University. The HPC Platform of ShanghaiTech University supported computation.

## Author contributions

J.Z. and Y.X. conceived the project and designed experiments. J.Z. performed organoid differentiation and characterization experiments and analysis of sequencing data. W.P. assisted with immunostaining and organoid differentiation. Y.L. and Y.Y. assisted with calcium imaging and MEA assay experiment. X.L. and J.W. assisted with DISC1 gene editing experiment. L.J., W.Z., Y.C., Q.Z., S.Chen, S.Chu, Z.J., D.H., Y.L., Y.T. and X.Z. contributed to cell culture. C.C. assisted with experimental resources. J.Z. and Y.X. prepared the manuscript.

## Competing interests

ShanghaiTech University has filed a patent application covering the generation and application of human serotonergic-enriched rostral and caudal hindbrain organoids, as well as related assembloids, for modeling serotonergic modulatory pathways.

**Extended Data Fig. 1.**
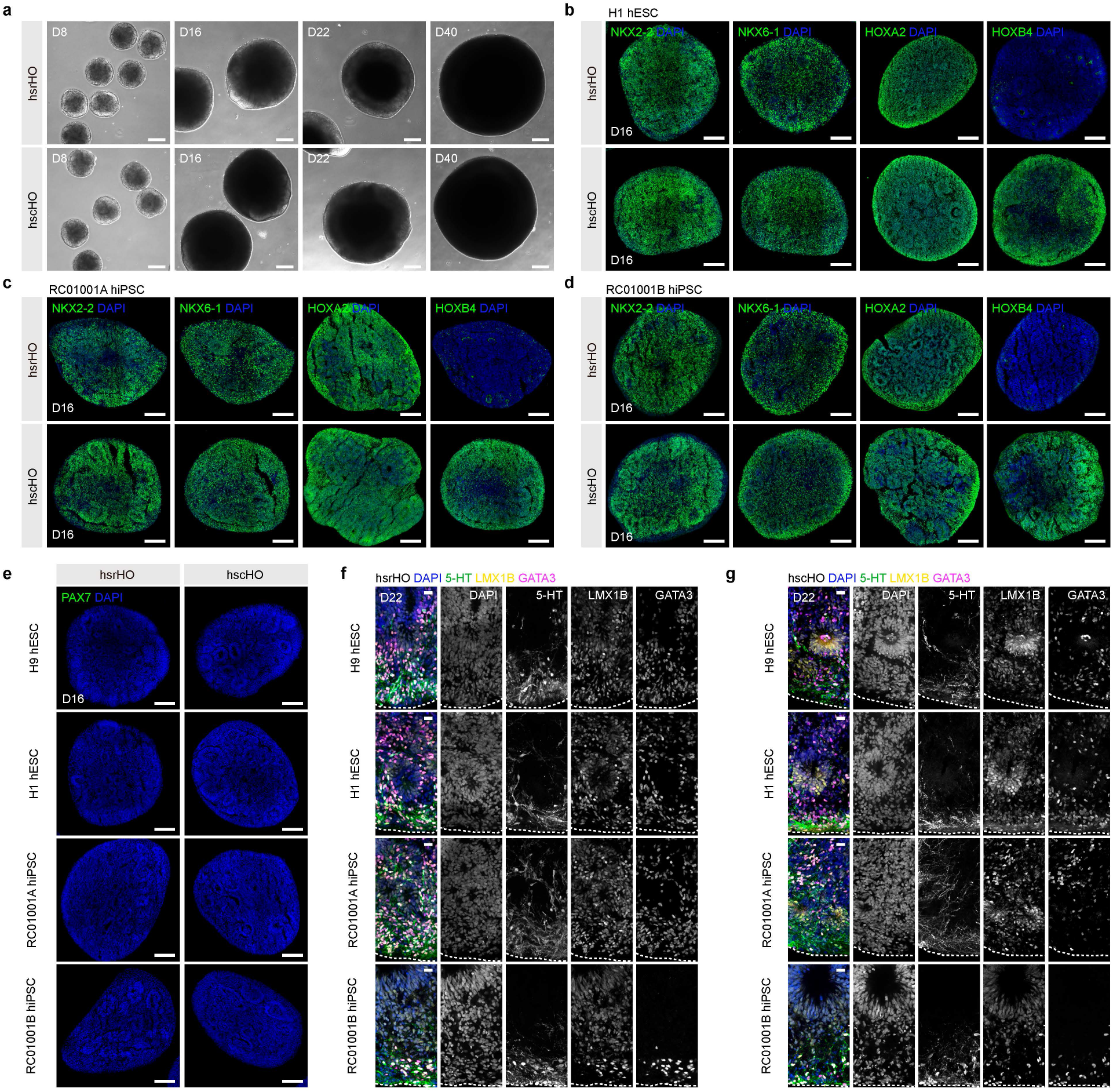
Generation of hsrHOs and hscHOs from hPSCs. **a**, Bright-field images of hsrHOs and hscHOs across development stages. **b-d**, Representative immunostaining images of NKX2-2, NKX6-1, HOXA2, and HOXB4 in day 16 hsrHOs and hscHOs generated from H1 hESC line (**b**), RC01001A hiPSC line (**c**) and RC01001B hiPSC line (**d**). **e**, Immunostaining of dorsal hindbrain marker (PAX7) of day 16 hsrHOs and hscHOs generated from hESC lines (H9 and H1) and hiPSC lines (RC01001A and RC01001B). **f-g**, Immunostaining of SN progenitor markers (LMX1B and GATA3) and 5-HT of day 22 hsrHOs (**f**) and hscHOs (**g**) generated from hESC lines (H9 and H1) and hiPSC lines (RC01001A and RC01001B). Scale bars, 200 μm (a, b, c, d, e), 50 μm (f, g).

**Extended Data Fig. 2.**
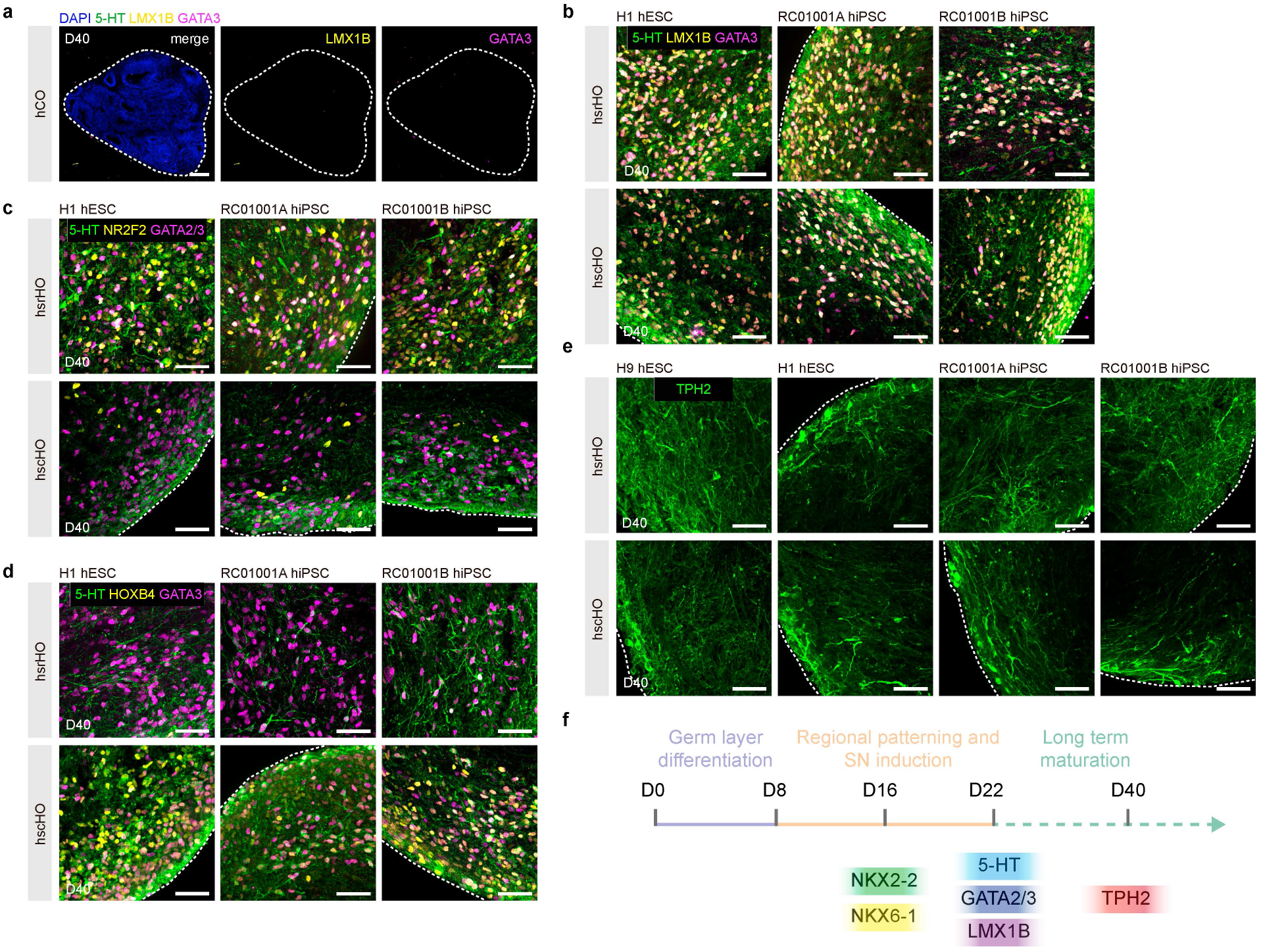
Characterization of SNs in hsrHOs and hscHOs. **a-b**, Immunostaining for SN markers (5-HT, LMX1B, and GATA3) of day 40 hCOs generated from H9 hESC line (**a**), and day 40 hsrHOs and hscHOs generated from hESC line (H1) and hiPSC lines (RC01001A and RC01001B) (**b**). **c-d**, Representative immunostaining images for rostral SN marker (5-HT, GATA2/3, and NR2F2) (**c**), and caudal SN marker (5-HT, GATA3, and HOXB4) (**d**) of day 40 hsrHOs and hscHOs generated from hESC line (H1) and hiPSC lines (RC01001A and RC01001B). **e**, Representative immunostaining images for mature SN marker (TPH2) of day 40 hsrHOs and hscHOs generated from hESC lines (H9 and H1) and hiPSC lines (RC01001A and RC01001B). **f**, Schematic diagram of hsrHOs and hscHOs development stages and the time when SN marker genes began to be expressed. Scale bars, 200 μm (a), 50 μm (b, c, d, e).

**Extended Data Fig. 3.**
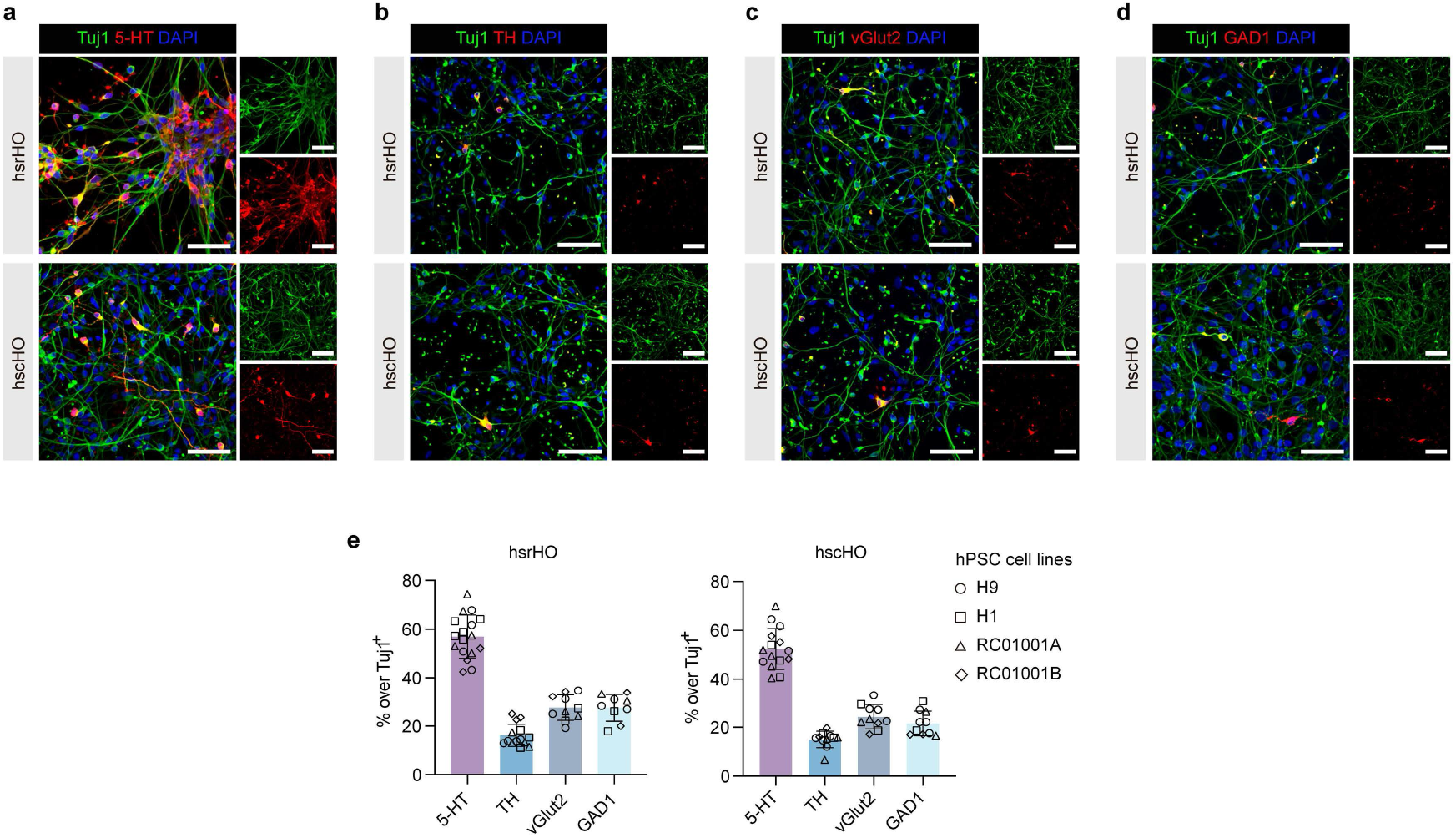
Characterization of neurons in hsrHOs and hscHOs. **a-e**, Representative immunostaining images for neuron marker (Tuj1) and SN marker (5-HT) (**a**), visceral motor neuron marker (TH) (**b**), glutamatergic neuron marker (vGlut2) (**c**) and GABAergic neuron marker (GAD1) (**d**) of day 40 dissociated hsrHOs and hscHOs, and quantification of each neuronal subtype among Tuj1-positive neuron (**e**) in day 40 dissociated hsrHOs and hscHOs generated from hESC lines (H9 and H1) and hiPSC lines (RC01001A and RC01001B) (hsrHO: n = 17 for 5-HT, n = 14 for TH, n = 10 for vGlut2, n = 9 for GAD1, one to four differentiations from four hPSC cell lines; hscHO: n = 15 for 5-HT, n = 11 for TH, n = 10 for vGlut2, n = 10 for GAD1, one to four differentiations from four hPSC cell lines). Scale bars, 200 μm (a, b, c, d).

**Extended Data Fig. 4.**
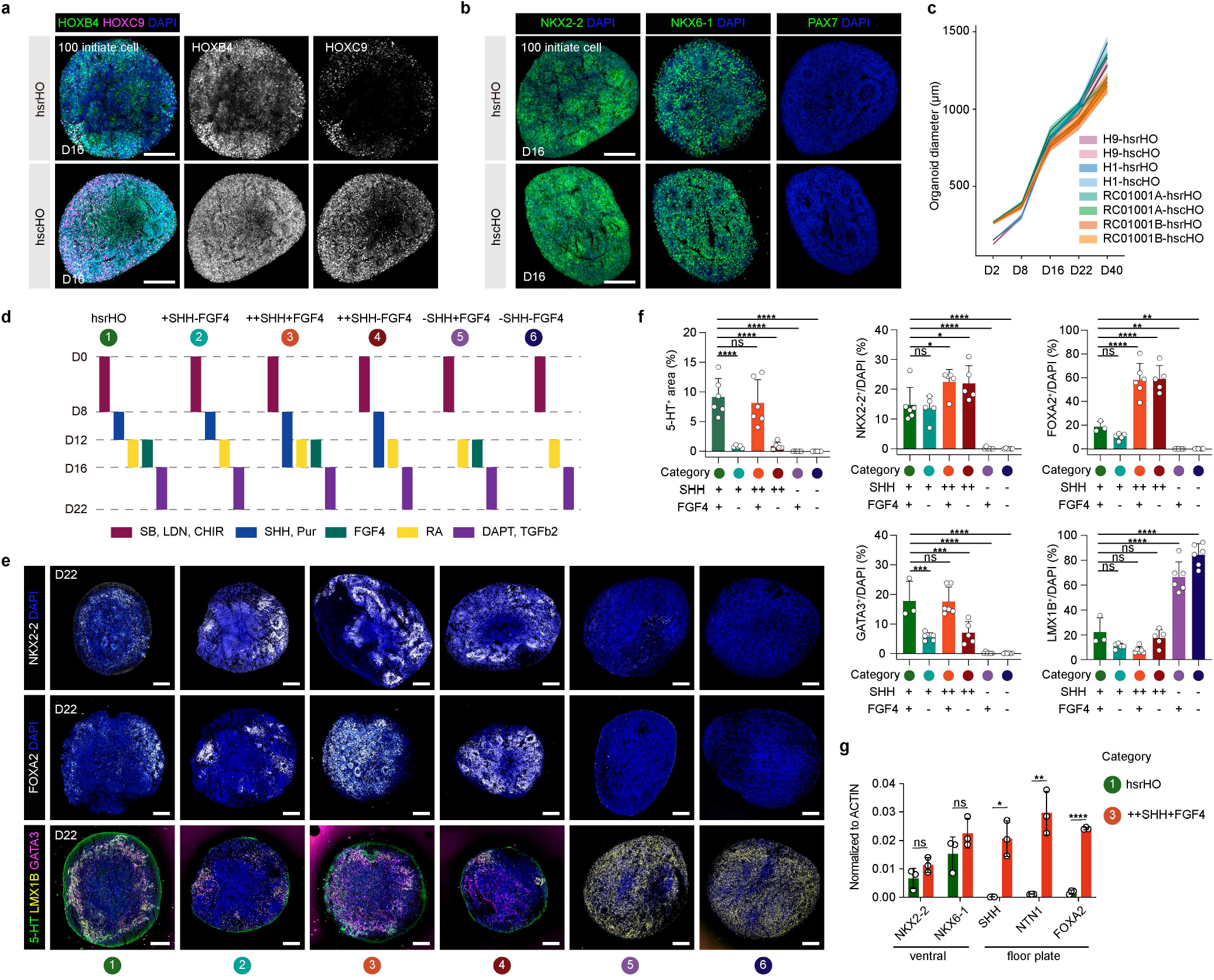
Factors affecting SNs patterning and generation. **a**, Representative immunostaining images of rostral-caudal markers in hsrHOs and hscHOs generated from H9 hESC cells seeded at low density (100 versus 500 cells). **b**, Representative immunostaining images of dorsal-ventral markers in hsrHOs and hscHOs generated from H9 hESC cells seeded at low density (100 versus 500 cells). **c**, Quantification of organoid size at different development stages (hsrHO: H9, n = 21, 27, 26, 27, 24 of day 2, 8, 16, 22, 40 from five differentiations, H1, n = 22, 28, 29, 27, 25 of day 2, 8, 16, 22, 40 from five differentiations, RC01001A, n = 22, 29, 28, 26, 28 of day 2, 8, 16, 22, 40 from five differentiations, RC01001B, n = 22, 29, 29, 24, 24 of day 2, 8, 16, 22, 40 from five differentiations; hscHO: H9, n = 21, 25, 26, 27, 23 of day 2, 8, 16, 22, 40 from five differentiations, H1, n = 20, 27, 28, 24, 28 of day 2, 8, 16, 22, 40 from five differentiations, RC01001A, n = 22, 28, 26, 26, 29 of day 2, 8, 16, 22, 40 from five differentiations, RC01001B, n = 21, 27, 29, 25, 27 of day 2, 8, 16, 22, 40 from five differentiations), data were presented as mean ± SEM. **d**, Schematic diagram describing the experimental design for assessing the effect of SHH and FGF4 on SNs differentiation. **e-f**, Representative immunostaining of ventral progenitor marker (NKX2-2), floor plate cell marker (FOXA2) and SN marker (5-HT, LMX1B and GATA3) across conditions (**e**), and quantification (**f**) (5-HT^+^ area, n = 6, 5, 6, 5, 6, 6 for category 1-6, two to three differentiations, ****p < 0.0001 for category 2 versus 1, p = 0.4225 for category 3 versus 1, ****p < 0.0001 for category 4 versus 1, ****p < 0.0001 for category 5 versus 1, ****p < 0.0001 for category 6 versus 1, ordinary one-way ANOVA; NKX2-2, n = 6, 5, 5, 5, 6, 6 for category 1-6, two to three differentiations, p = 0.6610 for category 2 versus 1, *p = 0.0121 for category 3 versus 1, *p = 0.0134 for category 4 versus 1, ****p < 0.0001 for category 5 versus 1, ****p < 0.0001 for category 6 versus 1, ordinary one-way ANOVA; FOXA2, n = 3, 5, 6, 5, 6, 6 for category 1-6, two to three differentiations, p = 0.3996 for category 2 versus 1, ****p < 0.0001 for category 3 versus 1, ****p < 0.0001 for category 4 versus 1, **p = 0.0091 for category 5 versus 1, **p = 0.0095 for category 6 versus 1, ordinary one-way ANOVA; GATA3, n = 3, 5, 6, 5, 6, 6 for category 1-6, two to three differentiations, ***p < 0.0001 for category 2 versus 1, p > 0.9999 for category 3 versus 1, ***p = 0.0006 for category 4 versus 1, ****p < 0.0001 for category 5 versus 1, ****p < 0.0001 for category 6 versus 1, ordinary one-way ANOVA; LMX1B, n = 3, 5, 6, 5, 6, 6 for category 1-6, two to three differentiations, p = 0.2024 for category 2 versus 1, p = 0.0569 for category 3 versus 1, p = 0.8373 for category 4 versus 1, ****p < 0.0001 for category 5 versus 1, ****p < 0.0001 for category 6 versus 1, ordinary one-way ANOVA). Note that LMX1B is also a dorsal progenitor marker of hindbrain. **g**, Gene expression by qPCR of ventral progenitor marker and floor plate cell marker of day 22 hsrHO and long-term SHH activation condition (category 1 and 3, n = 3 differentiations, NKX2-2, p = 0.1427, NKX6-1, p = 0.1887, SHH, *p = 0.0294, NTN1, *p = 0.0230, FOXA2, ****p < 0.0001). Scale bars, 200 μm (a, b, e).

**Extended Data Fig. 5.**
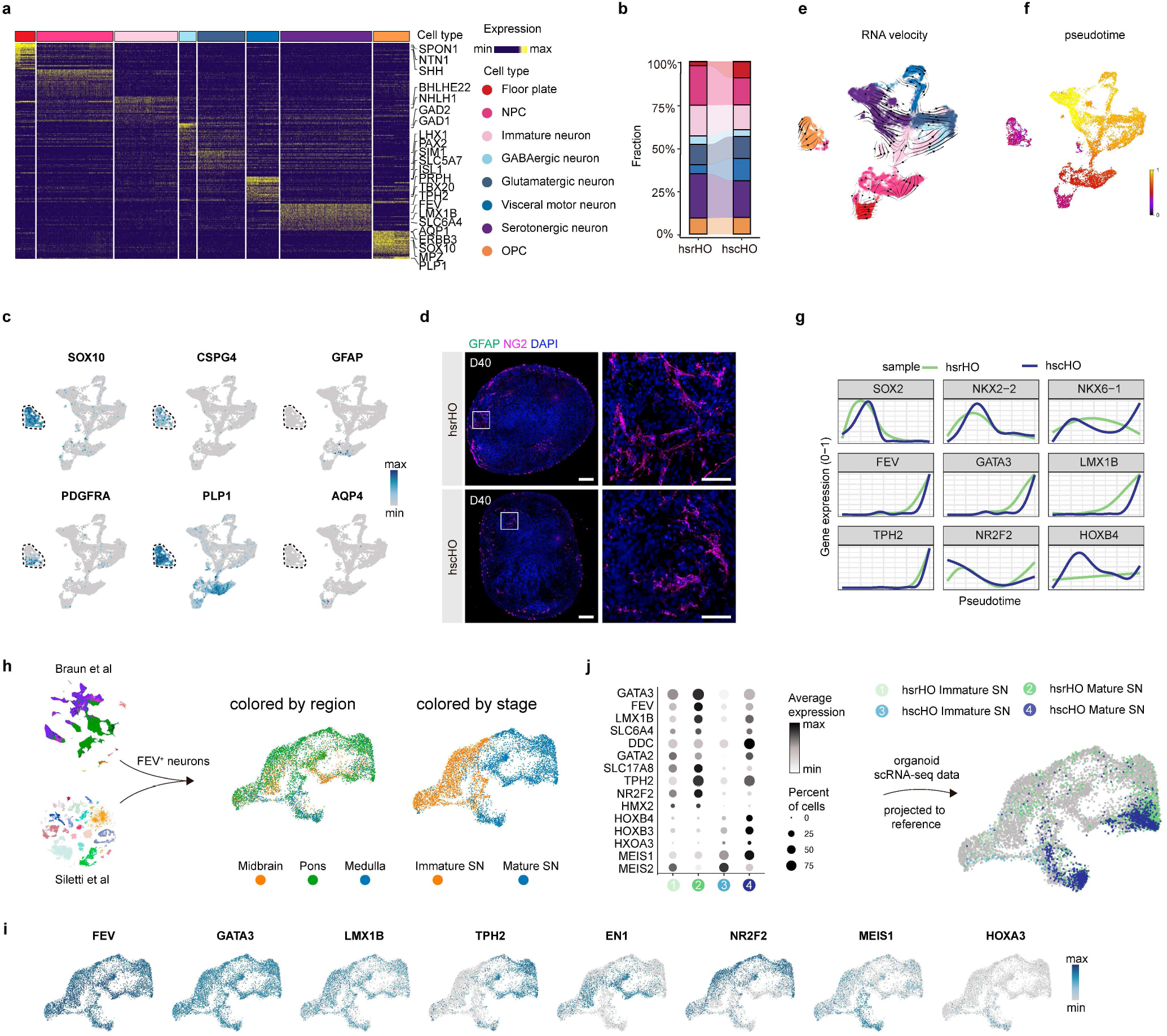
Molecular features of hsrHOs and hscHOs. **a**, Heatmap showing gene expression pattern of identified cell types in hsrHOs and hscHOs. **b**, Cell type proportions in hsrHOs and hscHOs. **c**, Feature plot showing the expression of OPC marker (SOX10, CSPG4, PDGFRA and PLP1) and astrocyte marker (GFAP and AQP4) in scRNA-seq profile. **d**, Representative immunostaining images for OPC marker CSPG4 (also known as NG2) and astrocyte marker GFAP. **e**, RNA velocity analysis of integrated hsrHOs and hscHOs scRNA-seq dataser. The stream arrows visualize the inferred trajectory. **f**, UMAP visualization of integrated hsrHOs and hscHOs scRNA-seq dataset, colored by pseudotime. **g**, Marker gene expression along pseudotime. **h**, Generation of a human SN reference by integrating SNs profiles from atlas published by Braun et al and Siletti et al. **i**, Feature plot showing the expression of marker genes in the human SN reference. **j**, Dot plot showing the expression of SN marker genes in hsrHOs and hscHOs and mapping organoid SN profile to human SN reference by label transfer. Scale bars, 200 μm (d), 50 μm (d(zoom)).

**Extended Data Fig. 6.**
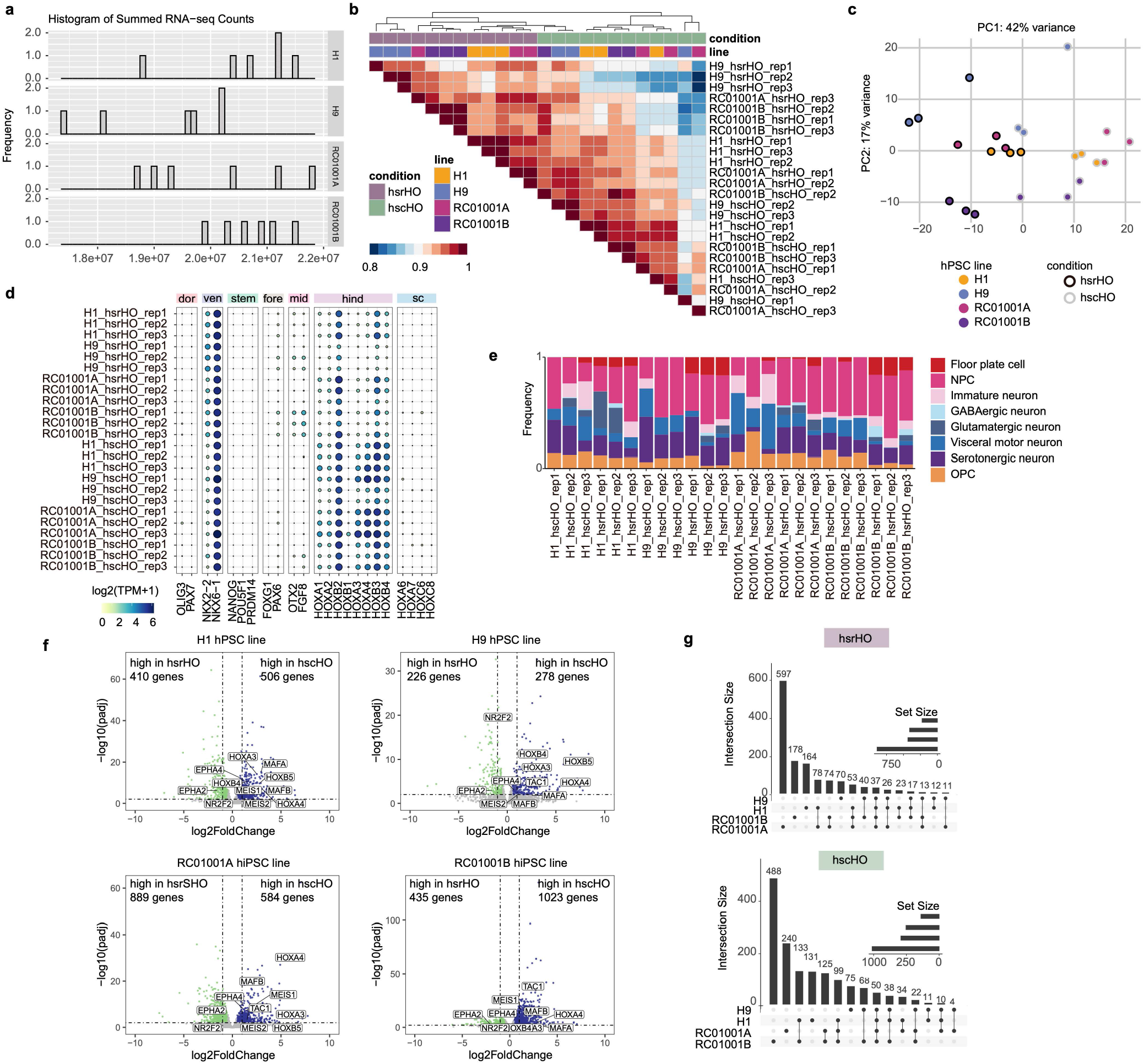
Reproducibility of hsrHOs and hscHOs generation. **a**, Histogram of counts for each bulk RNA-seq sample. **b**, Heatmap showing the spearman correlation between RNA-seq samples. **c**, PCA plot of RNA-seq samples. **d**, Dot plot showing the expression of marker genes. Dor, dorsal; ven, ventral; stem, stem cell; fore, forebrain; mid, midbrain; hind, hindbrain; sc, spinal cord. **e**, Cell proportions deconvolved using DWLS on bulk RNA-seq sample. **f**, Volcano plot showing the DEGs between hsrHOs and hscHOs generated from hESC lines (H9 and H1) and hiPSC lines (RC01001A and RC01001B). **g**, Intersection of genes up-regulated in hsrHOs (up) or hscHOs (bottom) generated from hESC lines (H9 and H1) and hiPSC lines (RC01001A and RC01001B).

**Extended Data Fig. 7.**
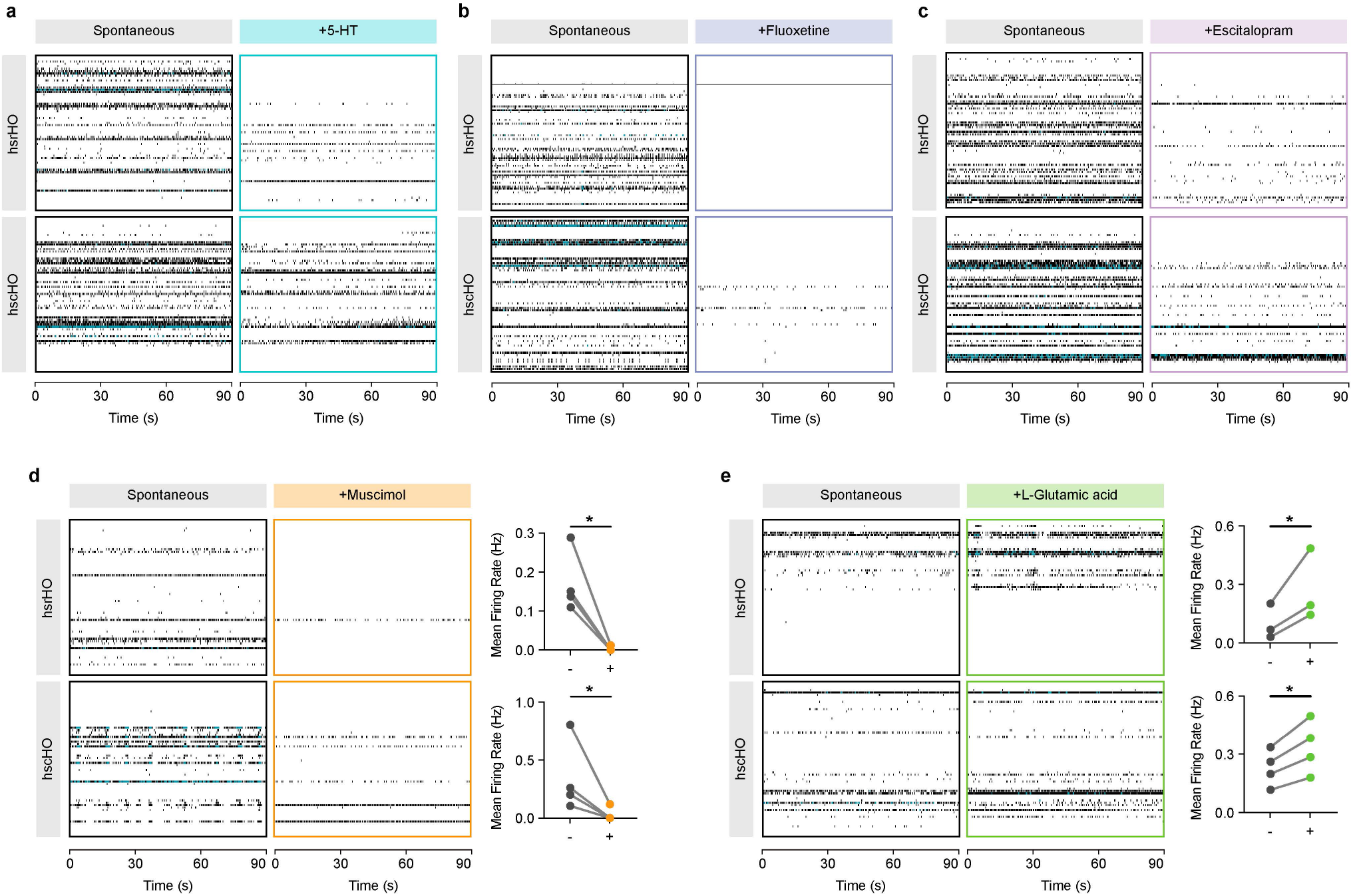
MEA assay of hsrHOs and hscHOs. **a-c**, Raster plots of MEA recordings of hsrHOs and hscHOs sections under spontaneous conditions and upon treatment with 5-HT (**a**), fluoxetine (**b**) or escitalopram (**c**). **d-e**, Raster plots and corresponding quantification of MEA recordings of hsrHOs and hscHOs sections under spontaneous conditions and upon treatment with muscimol (**d**) or L-glutamic acid (**e**) (For muscimol: n = 4 of hsrHOs and n = 4 for hscHOs, two differentiations, *p = 0.0109 for hsrHOs and *p = 0.0178 for hscHOs, ratio paired t test; for L-Glutamic acid: n = 3 of hsrHOs and n = 4 for hscHOs, two differentiations, *p = 0.0283 for hsrHOs, ratio paired t test, *p = 0.0151 for hscHOs, paired t test).

**Extended Data Fig. 8.**
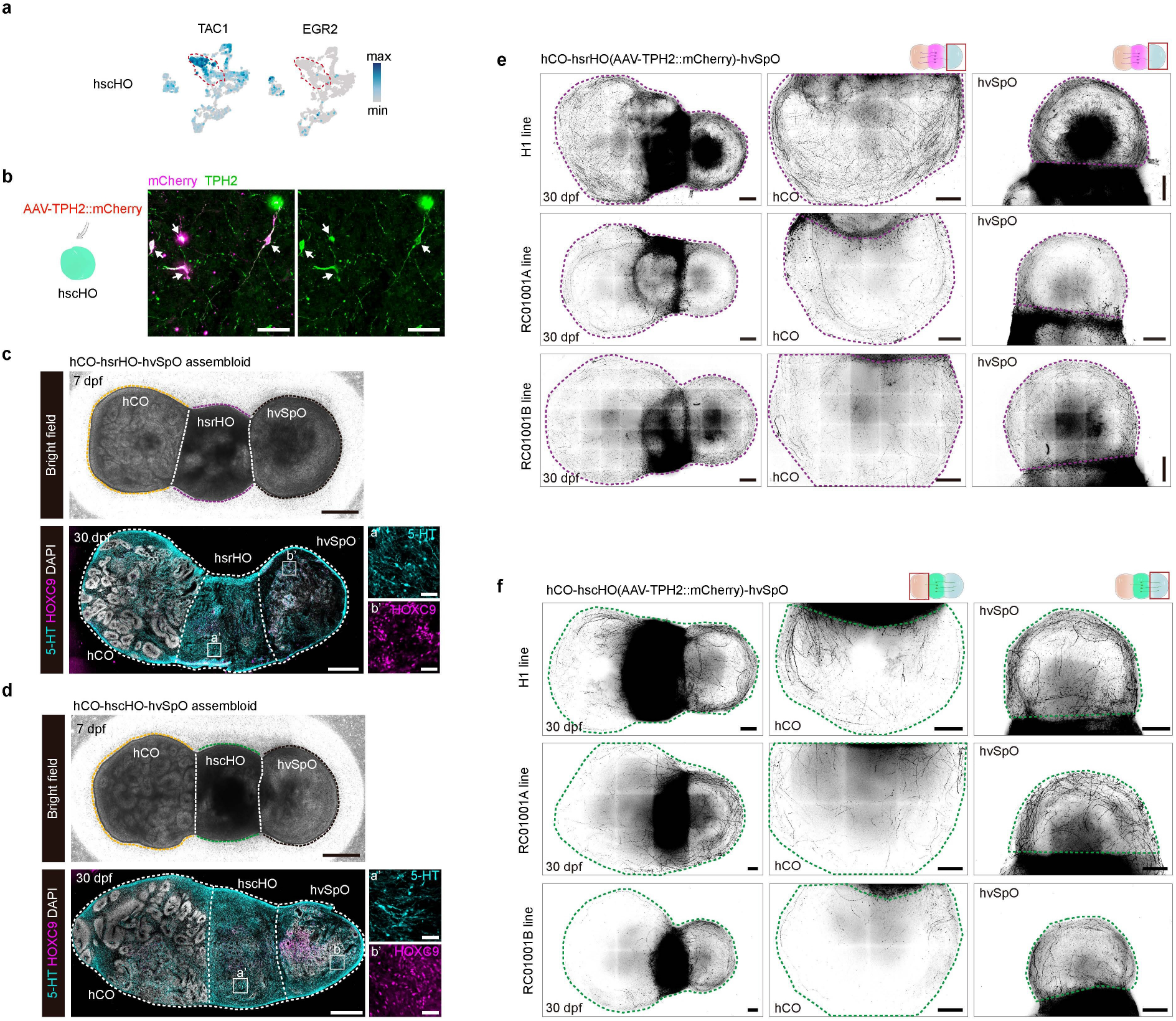
Serotonergic axon projections in assembloids. **a**, Feature plot of marker gene expression in day 40 hscHO scRNA-seq profile. **b**, Immunostaining of TPH2 and mCherry in day 60 hscHO infected with AAV9-TPH2::mCherry vectors. **c-d**, Bright-field and immunostaining for 5-HT and HOXC9 in hCO-hsrHO-hvSpO (**c**) and hCO-hscHO-hvSpO (**d**). **e-f**, Imaging of serotonergic axon projections (mCherry^+^) in hCO-hsrHO-hvSpO (**e)** and hCO-hscHO-hvSpO (**f**) assembloids. Scale bars, 500 μm (c, d, e, f), 50 μm (b, c(zoom), d(zoom)).

**Extended Data Fig. 9.**
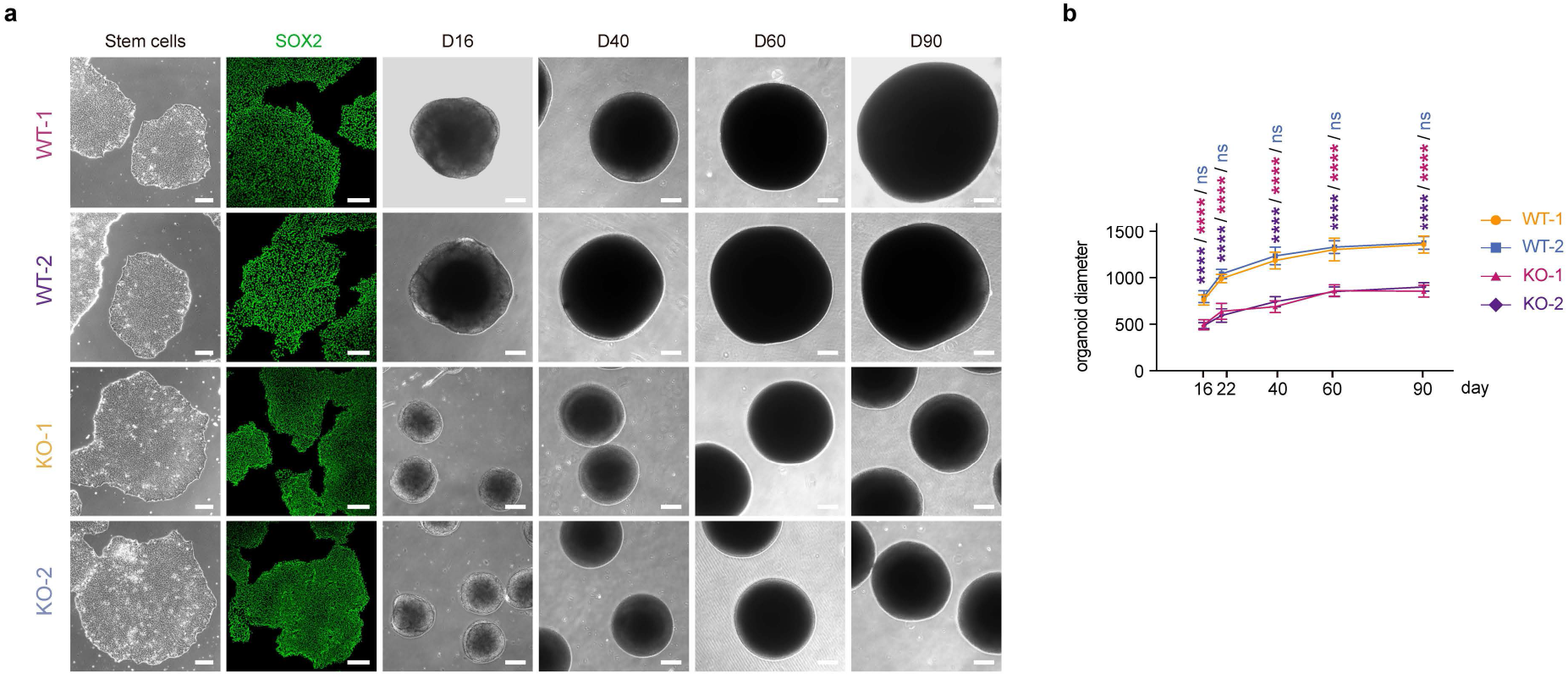
DISC1 deficiency in hscHOs. **a**, Bright-field images and SOX2 immunostaining of WT and DISC1 KO hESCs and bright-field images of hscHOs generated from WT and DISC1 KO hESCs. **b**, Quantification of the hscHO diameter across days in culture (WT-1, n = 12, 11, 12, 9, 7 of day 16, 22, 40, 60, 90, three differentiations; WT-2, n = 14, 11, 15, 10, 8 of day 16, 22, 40, 60, 90, three differentiations; KO-1, n = 15, 11, 18, 14, 16 of day 16, 22, 40, 60, 90, three differentiations; KO-2, n = 16, 12, 17, 14, 14 of day 16, 22, 40, 60, 90, three differentiations; day 16, p = 0.3392 for WT-2 versus WT-1, ****p < 0.0001 for KO-1 versus WT-1, ****p < 0.0001 for KO-2 versus WT-1; day 22, p = 0.2000 for WT-2 versus WT-1, ****p < 0.0001 for KO-1 versus WT-1, ****p < 0.0001 for KO-2 versus WT-1; day 40, p = 0.1868 for WT-2 versus WT-1, ****p < 0.0001 for KO-1 versus WT-1, ****p < 0.0001 for KO-2 versus WT-1; day 60, p = 0.6790 for WT-2 versus WT-1, ****p < 0.0001 for KO-1 versus WT-1, ****p < 0.0001 for KO-2 versus WT-1; day 90, p = 0.9257 for WT-2 versus WT-1, ****p < 0.0001 for KO-1 versus WT-1, ****p < 0.0001 for KO-2 versus WT-1, two-way ANOVA). Scale bars, 200 μm (a).

**Extended Data Table 1:**
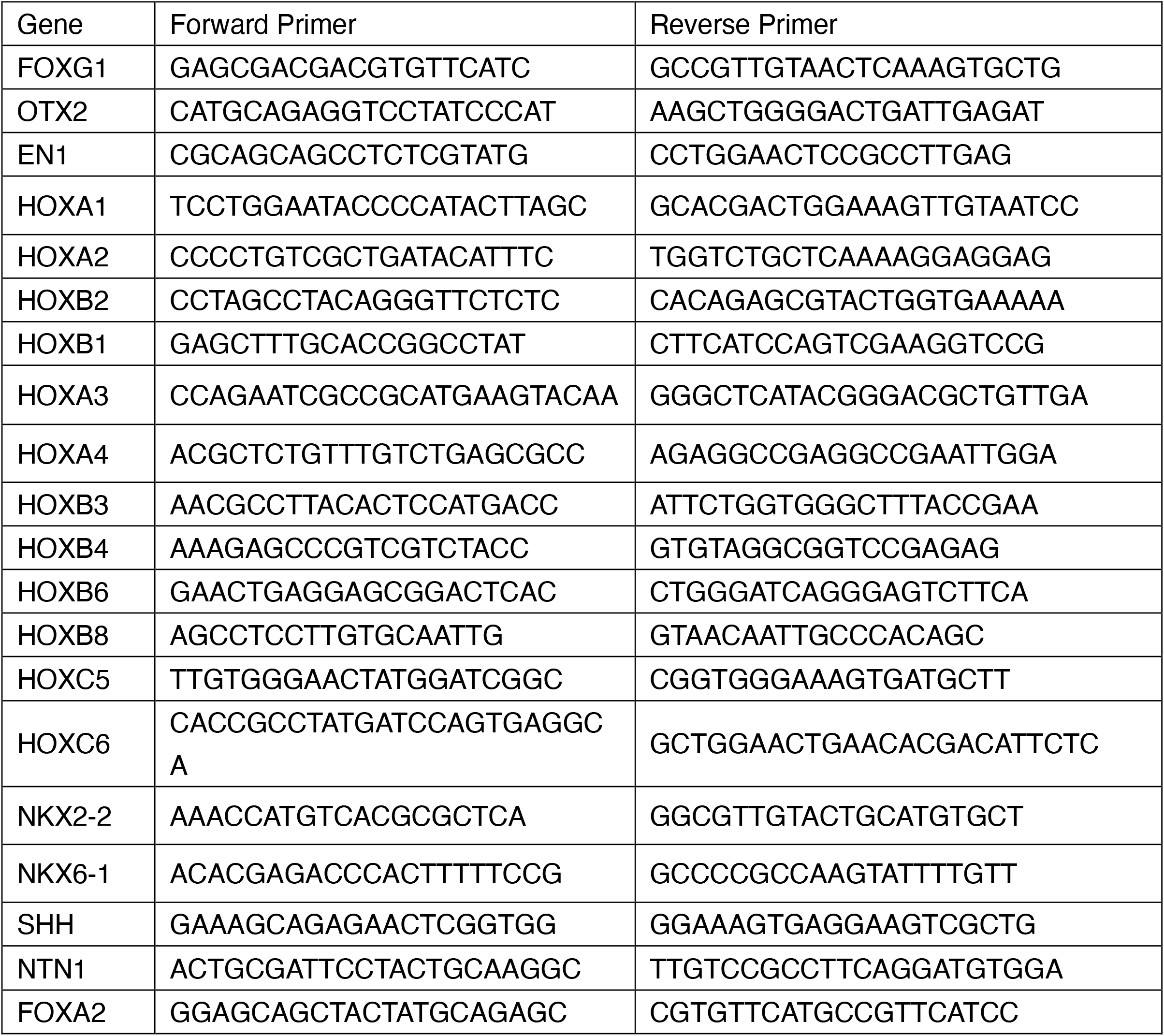
Primers used for qPCR experiments.

